# Osteoblast cell death triggers a pro-osteogenic inflammatory response regulated by reactive oxygen species and glucocorticoid signaling in zebrafish

**DOI:** 10.1101/2021.05.08.443237

**Authors:** Karina Geurtzen, Ankita Duseja, Franziska Knopf

**Affiliations:** CRTD - Center for Regenerative Therapies TU Dresden, Center for Healthy Aging TU Dresden, Germany; Department of Oncology and Metabolism, Metabolic Bone Centre, Sorby Wing Northern General Hospital, Sheffield, UK

**Keywords:** Zebrafish, osteoblast, macrophage, ablation, glucocorticoid, reactive oxygen species, lineage tracing

## Abstract

In zebrafish, transgenic labeling approaches, robust regenerative responses and excellent *in vivo* imaging conditions enable precise characterization of immune cell behavior in response to injury. Here, we monitored osteoblast-immune cell interactions in bone, a tissue which is particularly difficult to *in vivo* image in tetrapod species. Ablation of individual osteoblasts leads to recruitment of neutrophils and macrophages in varying numbers, depending on the extent of the initial insult, and initiates generation of *cathepsinK*+ osteoclasts from macrophages. Induced osteoblast death triggers the production of pro-inflammatory cytokines and reactive oxygen species, which are needed for successful macrophage recruitment. Excess glucocorticoid signaling as it occurs during the stress response inhibits macrophage recruitment, maximum speed and changes the macrophages’ phenotype. While osteoblast loss is compensated for within a day by contribution of committed osteoblasts, macrophages continue to populate the region. Their presence is required for osteoblasts to fill the lesion site. Our model enables visualization of homeostatic bone repair after microlesions at single cell resolution and demonstrates a pro-osteogenic function of tissue-resident macrophages in non-mammalian vertebrates.

**Summary statement:** Laser-mediated osteoblast ablation induces recruitment of tissue-resident macrophages by a release of reactive oxygen species. The presence of macrophages is required for osteoblasts to repopulate the lesion site and can be modulated by glucocorticoids.

## Introduction

The skeleton and the immune system are close interaction partners, and crosstalk between both, which is controlled by a set of regulatory molecules (Takayanagi, 2007), influences bone formation and affects bone regeneration. Excessive and prolonged activation of inflammatory cells causes bone destructive diseases such as rheumatoid arthritis, while long term treatment with anti-inflammatory steroids causes osteoporosis (X. Feng & McDonald, 2011; Takayanagi, 2007). These diseases are associated with pain and fragile bone, and represent major health issues with strongly increasing incidence in the aging population (Odén et al., 2015).

After injury, recruitment of immune cells is the first step to ensure proper healing and to prevent the spread of inflammation (Duffield, 2003). Neutrophils dominate the early inflammatory response, becoming attracted to the respective sites immediately after the insult, in order to clear debris and recruit macrophages (Kolaczkowska & Kubes, 2013). This recruitment is initiated by various cytokine and chemokine stimuli released by neutrophils and apoptotic cells at the wound site (Duffield, 2003). Early arriving macrophages display an inflammatory phenotype and release cytokines to induce tissue degradation and cell apoptosis (Diez-Roux & Lang, 1997; Leibovich & Ross, 1975). Resolution of inflammation during later wound healing and tissue repair is promoted by anti-inflammatory macrophages (Novak & Koh, 2013). In the bone fracture environment, macrophage contribution is essential for deposition and mineralization of bone matrix (Andrew et al., 1994). In particular, macrophages initiate bone remodeling by direct interaction with osteoblasts and osteoclasts in damaged bone (Batoon et al., 2017; Jilka et al., 2007). Moreover, macrophages produce osteoactive molecules which promote osteogenic differentiation and mineralization (Pettit et al., 2008; Sinder et al., 2015). Conversely, interaction with osteoblasts can induce cells of the monocyte/macrophage lineage to differentiate towards osteoclasts (Quinn et al., 1998).

Zebrafish has emerged as a powerful animal model to study immunity and inflammation (C. Hall et al., 2009; Niethammer et al., 2009; Renshaw et al., 2006; Trede et al., 2004), bone metabolism, remodeling and skeletal disease (Banerji et al., 2016; Carvalho et al., 2017; Hayes et al., 2013; Kimmel et al., 2010; McNulty et al., 2012; Witten & Huysseune, 2009). Skeletal and immune cell biology are largely conserved among vertebrates, and zebrafish share the respective involved cell types, signaling pathways and molecules with mammals (Renshaw & Trede, 2012; Witten & Huysseune, 2009). Compared to classic vertebrate models such as rodents (Brittijn et al., 2009) zebrafish research benefits from early and rapid bone development in the presence of optical transparency up to a late larval stage (Cubbage & Mabee, 1996). *In vivo* imaging of immune and skeletal tissue can be performed using a variety of transgenic tools labeling specific bone and immune cell types, enabling the visualization of cellular interactions in real time (Chen & Zon, 2009; Hammond & Moro, 2012).

Studies investigating the cellular reaction of zebrafish bone cells to injury have focused on the adult fin, in particular after amputation or cryoinjury (Ando et al., 2017; Chassot et al., 2016; Geurtzen et al., 2014; Knopf et al., 2011; Singh et al., 2012; Sousa et al., 2011), or on the zebrafish jaw (Ohgo et al., 2019; Paul et al., 2016; H. Zhang et al., 2015). During fin and scale regeneration, live imaging of injury-responsive osteoblasts identified their ability to migrate and dedifferentiate, but also revealed the importance of *de novo* osteoblast generation (Ando et al., 2017; Cox et al., 2018; Geurtzen et al., 2014). Larval zebrafish models have been employed to understand vertebrate bone development (Ahi et al., 2016; DeLaurier et al., 2019; Kimmel et al., 2010; Sharif et al., 2014; Tarasco et al., 2017) and to decipher pathomechanisms underlying congenital skeletal disease (Fiedler et al., 2018; Gistelinck et al., 2018; Tonelli et al., 2020). While *in vivo* imaging studies on immune cell recruitment after infection and injury of non-osseous tissues (axonal tissue, mesenchymal fin fold tissue) are widely used (Ellett et al., 2011; Tomoya Hasegawa et al., 2017; Isles et al., 2019; Li et al., 2012; Lieschke et al., 2001; Sanderson et al., 2015) sterile larval bone injury models are missing.

In this study, we present a novel laser-induced lesion paradigm in a developing skull bone in zebrafish, which provides a powerful tool to study the interaction between bone and immune cells *in vivo*. Using this model, we demonstrate the variable extent of immune cell recruitment in response to ablation of osteoblasts, illustrate the ablation-induced release of reactive oxygen species (ROS) and show that neutrophils, tissue-resident macrophages and *cathepsinK*+ osteoclast-like cells are attracted to dying osteoblasts, which are replenished by proliferation and migration of *osterix*+ osteoblasts. Macrophage recruitment is inhibited by the systemic application of antioxidants as well as glucocorticoid administration, which additionally changes macrophage phenotype. Ablation of macrophages by a nitroreductase-mediated approach leads to a reduction of osteoblasts at the lesion site. Our model can be used to elucidate the signals driving appropriate and disturbed macrophage and neutrophil recruitment to injured bone tissue *in vivo*, which is relevant for a variety of inflammatory bone diseases and for bone cell turnover during tissue homeostasis.

## Results

### A 10 % ablation of opercular osteoblasts is quickly reversed and leaves opercular growth unaffected

UV laser mediated cell ablations, which lead to loss of fluorescent signal produced by transgenic fluorophore reporters (Morsch et al., 2017), are known to effectively kill target cells in zebrafish (Dehnisch Ellström et al., 2019; Mathias et al., 2006; Smutny et al., 2015). In order to create a confined lesion in bone and simulate osteoblast cell death, we performed osteoblast ablation in transgenic *osterix*:nGFP zebrafish larvae at 6 days post fertilization (dpf), in which osteoblasts of the forming gill cover (opercle) are labeled by GFP, by using specific UV laser settings at a spinning disk confocal microscope. We evaluated the cell damage performed by laser ablation by investigating the number of opercular osteoblasts with and without lesion at several time points. At 1 hour post lesion (hpl) we detected a prominent loss of GFP signal at the lesion site (**Fig. 1A**), which corresponded to a loss of approximately 10 % opercular osteoblasts (100 +/− 9,9 cells in uninjured vs. 87,4 +/− 10,2 cells in lesioned zebrafish, **Fig. 1B**). At 1 day post lesion (dpl), recovery of GFP fluorescence in the lesion site was observed, despite the fact that osteoblast numbers remained slightly (but not significantly) lower than in control fish (103,4 +/− 7,3 cells in uninjured vs. 93,5 +/− 10,1 cells in lesioned zebrafish). Complete recovery of osteoblast number was achieved at 2 dpl (110,9 +/− 10,6 cells in uninjured vs. 106,1 +/− 7,9 cells in lesioned zebrafish, **Fig. 1B**), illustrating the quick recovery of osteoblast numbers in laser ablated opercles.

**Fig. 1:**
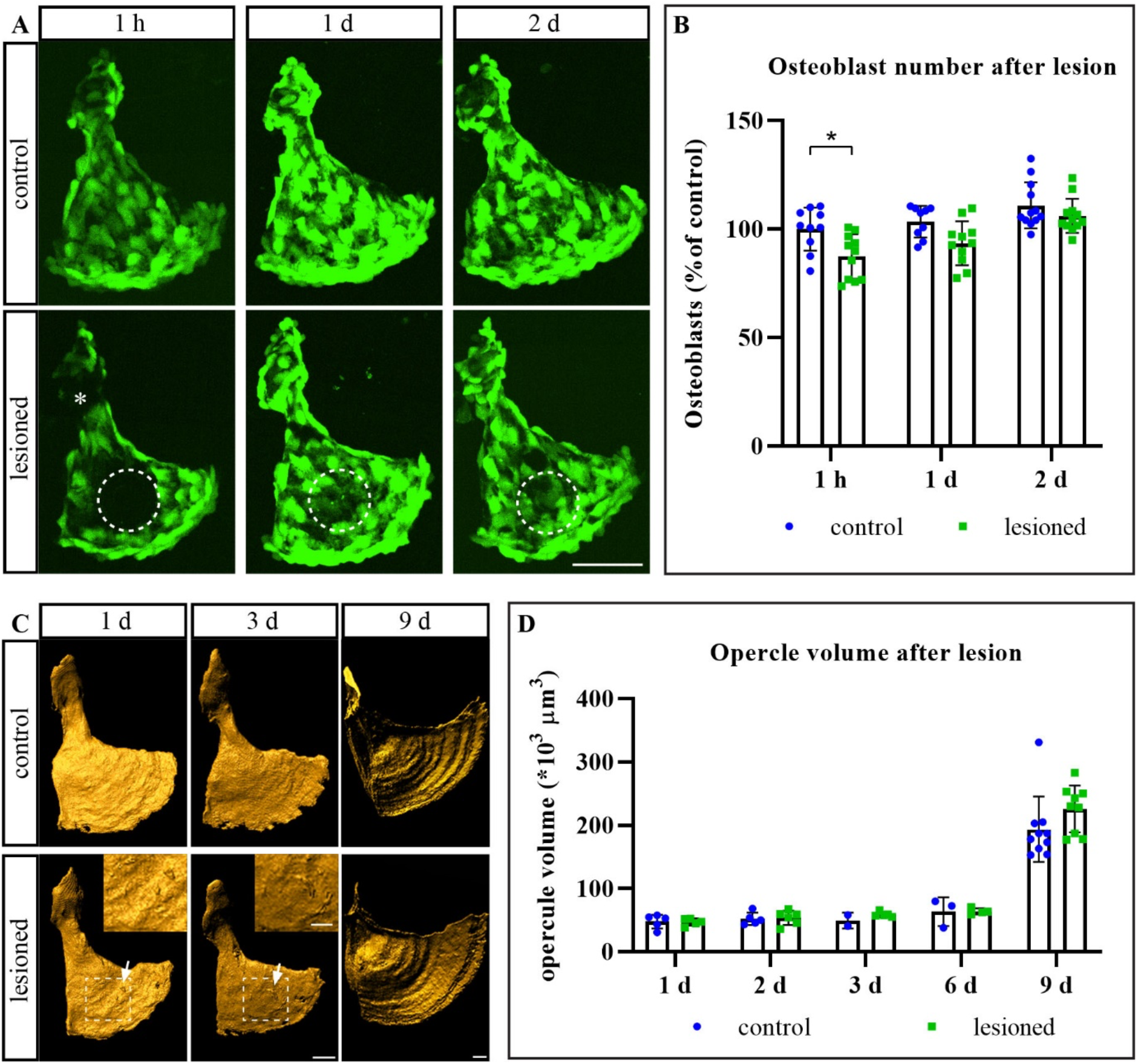
Osteoblast recovery and unaffected opercular growth after laser ablation. **A**: Representative images of the opercle region in transgenic *osterix*:nGFP 6 dpf larval zebrafish. Loss of osteoblasts at 1 hpl and full recovery at 2 dpl can be observed. White dashed line = ablated area. Scale bar = 50 µm. **B**: Quantification of osteoblast numbers from experiment shown in A. About 10 % of the osteoblasts are ablated. Mean + s.d. Students-t-test: * p = 0.011. n = 9-12. **C**: Opercles of 6 dpf lesioned zebrafish stained with alizarin red. Boxed area with arrows and inset: Laser traces in the form of two spaced rings. Scale bar overview = 20 µm. Scale bar insets = 10 µm. **D**: Quantification of opercular volume from experiment shown in C. No significant differences can be observed. Mean + s.d. Sidaks’ ANOVA. n = 3-10. h = hours, d = days.

Next, we characterized the effect of laser-assisted osteoblast lesions on opercle structure and growth. To evaluate opercle volume before and after lesion, we stained zebrafish larvae by alizarin red live staining, which labels calcified structures (Javidan & Schilling, 2004), and rendered the surface with the help of IMARIS software. Ablation of osteoblasts led to a distinct structural change of the calcified matrix in the form of two closely spaced rings in places where the laser had hit (arrow and insets in **Fig. 1C**). These marks could be observed for several days post lesion, and disappeared by 9 dpl (**Fig. 1C**). As these marks were quite prominent, and because bone forming osteoblasts were ablated, we wondered whether a change in opercle volume would result from lesion. Notably, quantification of opercular volume across different stages showed that there were no significant differences between lesioned and respective control zebrafish larvae (uninjured vs lesioned, all *10^3^ µm^3^, 1 dpl: 47,7 +/− 11,0 vs. 47,0 +/− 5,6, 2 dpl: 52,08+/−9,9 vs. 53,4 +/− 11,1, 3 dpl: 49,4 +/− 12,5 vs. 58,9 +/− 4,1, 6dpl: 63,3 +/− 22,4 vs. 63,9 +/− 4,8, 9 dpl: 193,4 +/− 51,6 vs. 225,4 +/− 37,2, **Fig. 1D**). This indicates that osteoblast ablation does not grossly affect opercular growth rate, although a temporal and spatially restricted structural damage in mineralized matrix could be observed, and that osteoblast numbers recover very quickly after ablation of a significant portion of osteoblasts.

### Osteoblast numbers recover by proliferation and migration of committed osterix+ osteoblasts

Because quick recovery of osteoblast numbers was observed after lesion, we wondered how this recovery was accomplished. The most plausible explanation to us seemed to be the possibility that remaining osteoblasts proliferated in order to restore the necessary pool of osteoblasts. To test this, we performed live imaging of transgenic *osterix*:nGFP x *histone*:mCherry zebrafish larvae, in which nuclei of osteoblasts can be identified by co-localization of GFP and mCherry (Knopf et al., 2011), and by using *osterix*:nGFP transgenic animals. Although osteoblast proliferation occurred (**Movie 1**), it was a rare event (observed in 1 out of 15 larvae). At the same time, we observed slow relocation of pre-existing osteoblasts, as indicated by an increased number of osteoblasts reaching into the lesion site at 12 hpl (**Fig. 2A**). To confirm migration of pre-existing osteoblasts into the ablation site, we performed CreERT2-loxP mediated lineage tracing of *osterix*+ osteoblasts. *osterix*:CreERT2-p2a-mCherry x *hsp70*:R2nlsG double transgenic fish (Geurtzen et al., 2014; Hans et al., 2009; Knopf et al., 2011) were either treated with 4-hydroxytamoxifen (4-OHT) to induce CreERT2 activity and excision of a loxP-flanked DsRed Stop cassette in osteoblasts, or the vehicle control one day before the lesion (**Fig. 2B**). Three days post osteoblast ablation the resulting nuclear GFP+ osteoblasts representing recombined cells and their progeny were visualized with the help of a heat shock (**Fig. 2B, C**) (Geurtzen et al., 2014; Hans et al., 2009; Knopf et al., 2011). In 4-OHT treated larvae, GFP+ osteoblasts accumulated at the lesion site (**Fig. 2C**, white arrow), while no GFP+ cells were detectable in the vehicle control. These results indicate that committed *osterix*+ opercular osteoblasts move into the ablation site to replenish the lost osteoblasts, a process which is likely supported by proliferation of the very same cells. However, future assays will be needed to test the possibility of *de novo* osteoblast formation from alternative sources.

**Fig. 2:**
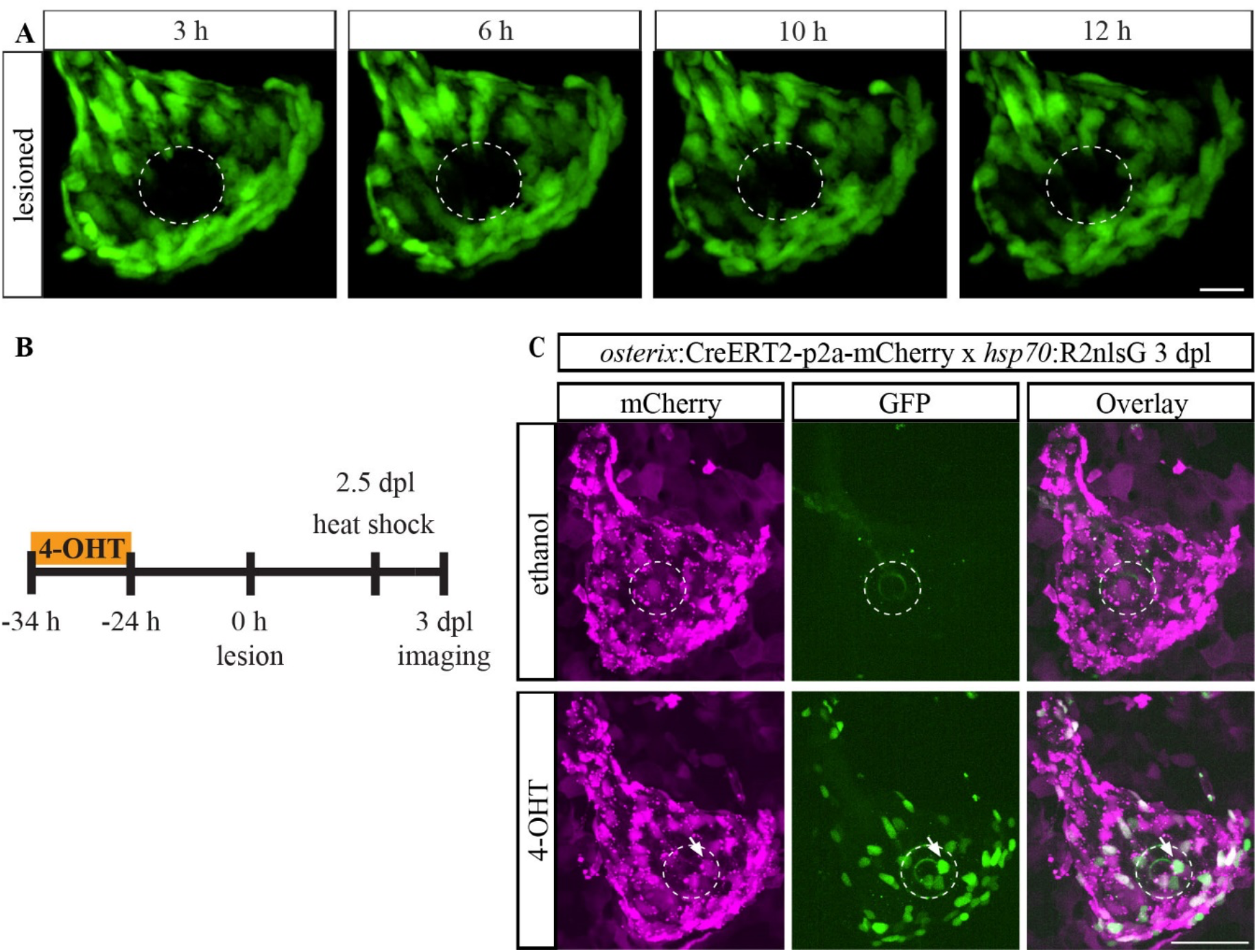
Pre-existing *osterix*+ osteoblasts migrate into the lesion site after ablation. **A**: Images of opercular osteoblasts in 6 dpf transgenic *osterix*:nGFP larval zebrafish. Cellular extensions reach into the lesion site (area within white dashed line) within several hpl. Scale bar = 20 µm. n = 3. **B**: Scheme on CreERT2-loxP mediated lineage tracing approach of osteoblasts. Osteoblast ablation was performed at 6 dpf/12 h post 4-OHT/vehicle treatment. Two days later, a single heat shock was used to visualize nuclear GFP expression. **C**: Representative images of 4-OHT and vehicle treated *osterix*:CreERT2-p2a-mCherry x *hsp70*:R2nlsG larval zebrafish at 3 dpl (9 dpf). Pre-existing committed opercular osteoblasts are located at the lesion site (white arrow and dashed line). Scale bar = 50 µm. n = 5-7.

### Osteoblast cell death leads to the release of immune cell attractants

Tissue damage and cell death lead to the release of a variety of chemokines and other cytokines, which have the potential to attract immune cells to the site of wounding (Duffield, 2003; Keightley et al., 2014). Furthermore, both processes enhance the expression of extracellular matrix (ECM) modifiers such as Matrix metalloproteinase 9 (Mmp9), a collagenase associated with inflammation which is found in wounded zebrafish (LeBert et al., 2015). We made use of transgenic *mmp9*:EGFP zebrafish (Ando et al., 2017) to test whether Mmp9 expression is induced in zebrafish bone tissue upon osteoblast ablation. While occasional GFP fluorescence was observed in the lesioned area at 1 dpl, we detected robust induction of GFP at the lesion site at 2 dpl (uninjured vs. lesioned, 1 hpl: 109,6 +/− 1,5 units vs. 108,6 +/− 0,3 units, 1 dpl: 112,2 +/− 8,3 units vs. 126,1 +/− 18,9 units, 2 dpl: 109,9 +/− 0,2 units vs. 124,3 +/− 7,0 units, **Fig. 3A, B**). We set out to identify earlier signs of inflammatory cues after osteoblast ablation and turned to reactive oxygen species (ROS), which are known to be produced soon after acute wounding of other tissues such as the larval fin fold, where they are responsible for leukocyte attraction to the site of injury (Niethammer et al., 2009), or the tail (Romero et al., 2018). In order to test whether ROS were generated after laser-assisted osteoblast ablation, we pre-soaked *osterix*:nGFP larval zebrafish in CellROX orange dye, which starts to fluoresce upon ROS presence, performed lesions and concomitant *in vivo* imaging. Almost instantaneous activation of ROS-caused fluorescence was detected within a minute after ablation, and lasted throughout the imaging period of approximately 20 minutes. In contrast, control larvae which had not been lesioned but equally soaked in the CellROX orange dye, did not show any signs of fluorescence (**Fig. 3C, Movies 2 and 3**). These results indicate that sterile, laser-assisted ablation of a low number of bone forming cells triggers a similar response to injury as seen in other, more severe injury paradigms such as tissue resection. They also hint at a potential ability of the lesion paradigm to trigger recruitment of immune cells and osteoclasts (Callaway & Jiang, 2015), which, consequently, would allow the *in vivo* observation of leukocyte interactions with osteoblasts in bone tissue.

**Fig. 3:**
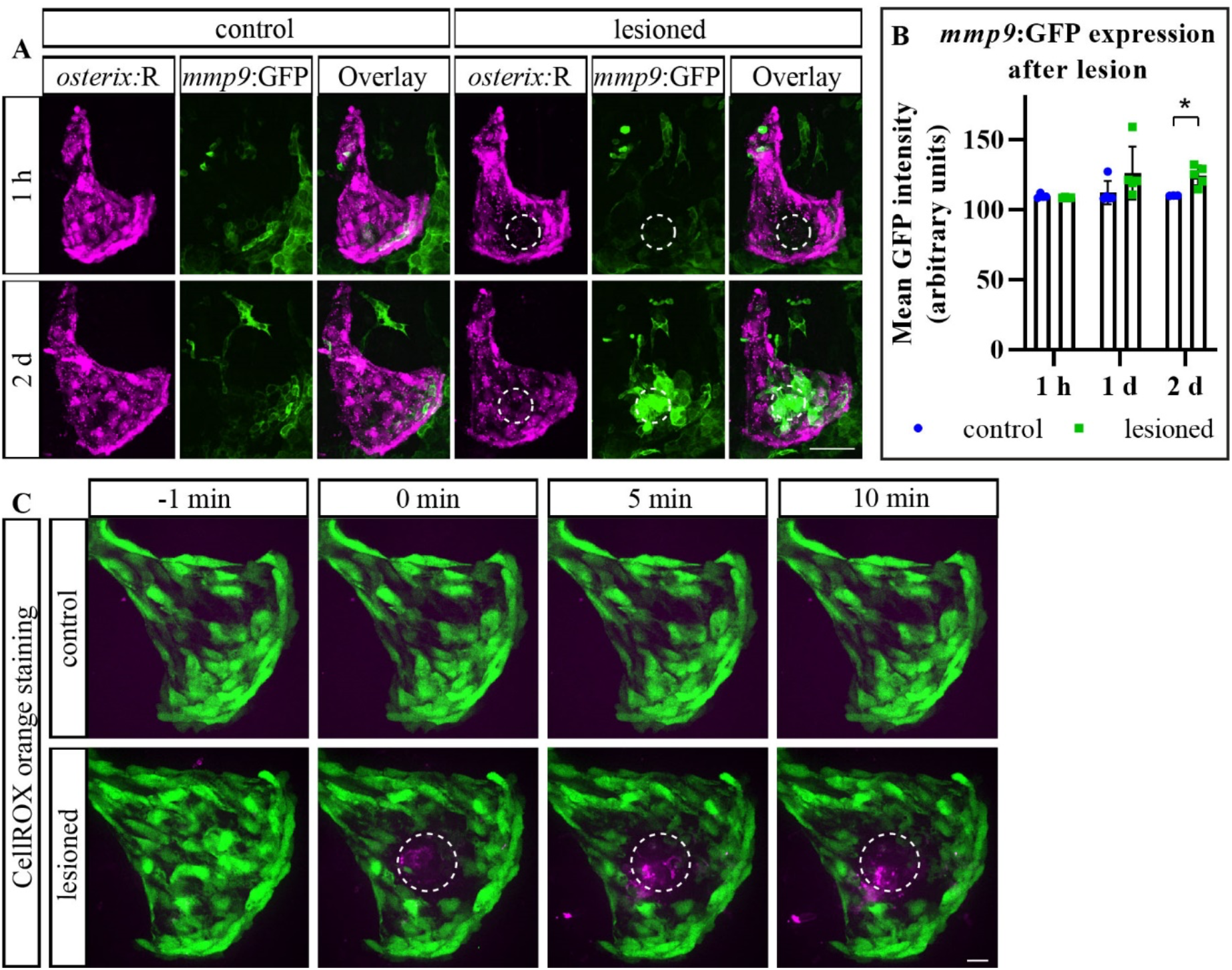
Laser-assisted osteoblast ablation triggers Mmp9 production and ROS release. **A**: Transgenic *osterix*:RFP x *mmp9*:GFP lesioned zebrafish showing robust *mmp9* activity at 2 dpl. White dashed line = lesioned area. Scale bar = 50 µm. **B**: Quantification of experiment shown in A. At 2 dpl GFP is significantly increased. Mean + s.d. Welch’s T-test: *p = 0.010. n = 3-5. **C**: CellRox orange staining of lesioned and unlesioned transgenic *osterix*:nGFP zebrafish larvae. ROS release is visible immediately after the laser lesion and increases over time in the ablation area (white dashed line). Scale bar = 10 µm. n = 6-7.

### Neutrophils, inflammatory macrophages and osteoclast-like cells become recruited to the lesion site

Cell death and the release of corresponding signals serve as triggers for recruitment of immune cells (Duffield, 2003; Keightley et al., 2014). Increased levels of *mmp9*:GFP expression and ROS after osteoblast ablation prompted us to test whether neutrophil numbers change upon lesion. Live imaging of double transgenic *osterix*:RFP x *mpo*:GFP zebrafish larvae labeling osteoblasts and neutrophils at the same time revealed fast recruitment of neutrophils into the lesion area within minutes (**Fig. 4A**, **Movie 4**).

**Fig. 4:**
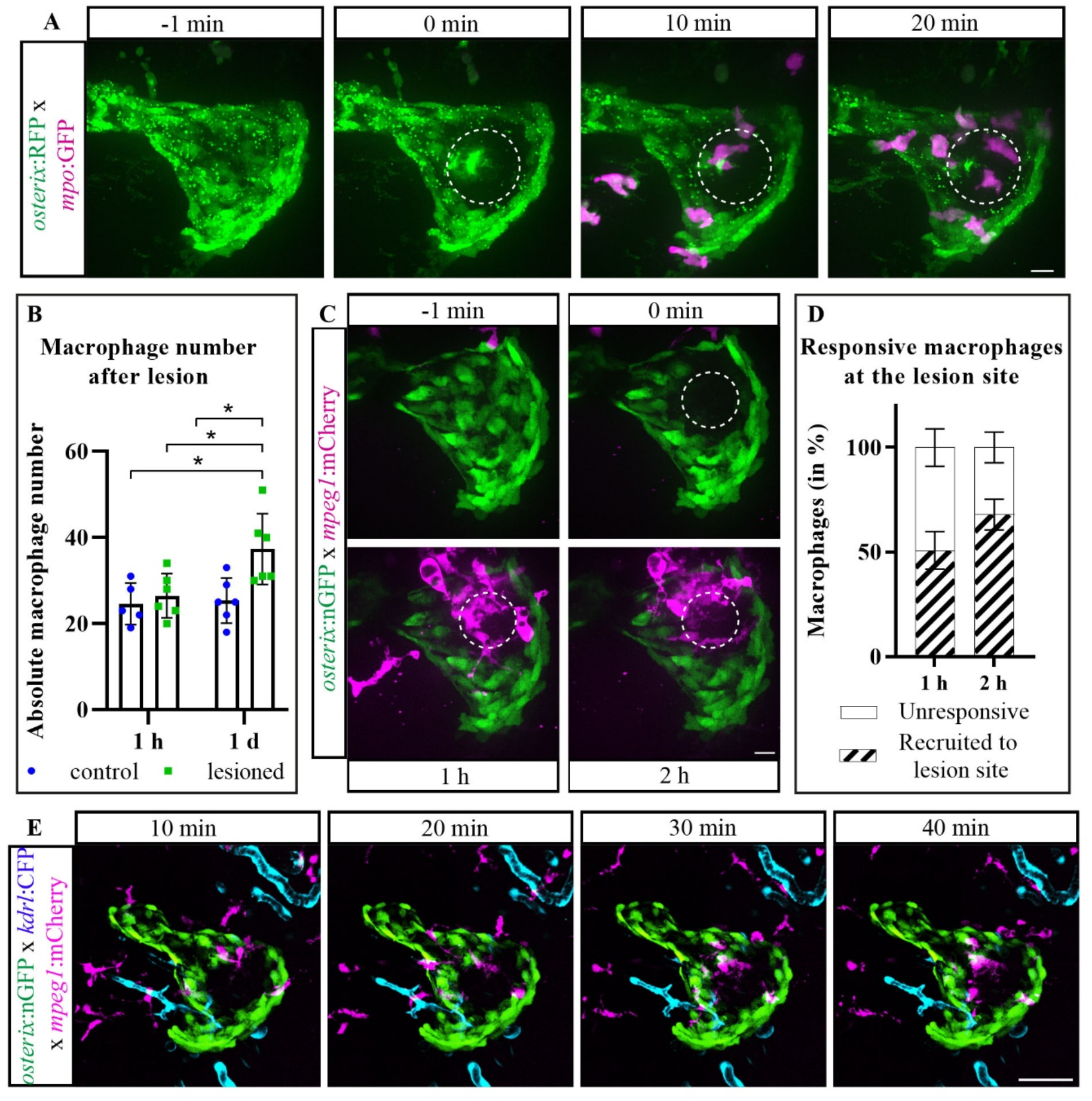
Neutrophil and macrophage recruitment to the site of osteoblast ablation. **A**: Time series of the opercle region in transgenic *osterix*:RFP x *mpo*:GFP zebrafish (RFP depicted in green). *mpo*:GFP+ neutrophils, depicted in magenta, migrate into the ablation site (white dashed line) within several minutes post ablation. Scale bar = 10 µm. n = 7. **B**: Quantification of macrophage number in the opercle area. The number increases significantly after 1 day in lesioned opercles. Mean + s.d. Tukey’s ANOVA: *p = 0.014 (1 d control vs. 1 d lesioned). n = 5-6. **C**: Time series of the opercle region of transgenic *osterix*:nGFP x *mpeg1*:mCherry zebrafish in which macrophages, depicted in magenta, migrate into the ablation site (white dashed line). Scale bar = 10 µm. **D**: Quantification of responsive macrophages migrating into the ablated area in the experiment shown in C. Mean + s.e.m. Sidak’s ANOVA. n = 5. E: Time series of the opercle region of transgenic *osterix*:nGFP x *kdrl*:CFP x *mpeg1*:mCherry zebrafish showing that macrophages, depicted in magenta, are recruited from the surrounding tissue and not from blood vessels (ablation site indicated by white dashed line). Scale bar = 50 µm. n = 3.

Similar to neutrophils, macrophages are attracted to the site of injury by cytokines and ROS (Mosser & Edwards, 2008). Moreover, early arriving neutrophils recruit macrophages to the injured or infected area after being the initial responders to the insult (Kolaczkowska & Kubes, 2013). We quantified the number of macrophages labeled by mCherry in double transgenic *osterix*:nGFP x *mpeg1*:mCherry zebrafish at different time points post lesion. While absolute macrophage numbers in the field of view did not change at 1 hpl, their number significantly increased until 1 dpl (uninjured vs. lesion, 1 hpl: 24,6 +/− 4,8 cells vs. 26,5 +/− 5,1 cells, 1 dpl: 25,3 +/− 5,2 cells vs. 37,3 +/− 8,3 cells, **Fig. 4B**), which suggests recruitment of macrophages to the lesion site after neutrophil arrival. Live-imaging using the above double transgenic zebrafish confirmed fast recruitment of macrophages that had resided in the field of view into the osteoblast-ablated area, as well of slightly delayed recruitment of macrophages from outside the field of view starting around 20 min (**Fig. 4C, Movie 5**). More than 50 % of macrophages passing the field of view during the imaging time were attracted into the lesion site during the first and second hour post lesion (1 hpl: 50,8 +/− 9,0 %, 2 hpl: 68,0 +/− 7,4 %, **Fig. 4D**). Live-imaging of *osterix*:nGFP x *mpeg1*:mCherry zebrafish combined with a transgenic marker for endothelial tissue, *kdrl*:CFP (Hess & Boehm, 2012), revealed that macrophages attracted to the lesion site arrive from within the tissues close to the lesion site and not from the blood stream, confirming their tissue-residency (**Fig. 4E, Movie 6**).

Different macrophage subtypes and polarization stages can be observed in response to injury and inflammation (Stout et al., 2005). Early inflammatory responses are often associated with the inflammatory type of macrophages (Duffield, 2003), also in zebrafish (Nguyen-Chi et al., 2015). Using triple transgenic *osterix*:nGFP x *mpeg1*:mCherry x *irg1*:EGFP zebrafish larvae in which activated macrophages (Sanderson et al., 2015) are labeled alongside osteoblasts, an increase in activated macrophages (mCherry/EGFP double+ migratory cells) was detected at 1 dpl (100,0 +/− 53,8 % in uninjured vs. 169,6 +/− 69,6 % in lesioned, white arrows in **Fig. 5A, B**). Similarly, the use of *tnf-α*:EGFP x *mpeg1*:mCherry transgenic zebrafish (Marjoram et al., 2015) demonstrated increased numbers of inflammatory macrophages at 1 and 2 dpl (uninjured vs lesioned, 1 hpl: 6,0 +/− 2,9 % vs. 5,6 +/− 1,7 %, 1 dpl: 9,4 +/− 5,5 % vs. 18,0 +/− 5,1 %, 2 dpl: 14,8 +/− 6,3 % vs. 24,2 +/− 9,4 %, white arrows in **Fig. 5C, D**). However, the majority of recruited macrophages at 1 and 2 dpl did not show the *tnf-α*+ inflammatory phenotype, which was only detected in about 20 % of all macrophages (1 dpl: 18,0 +/− 5,1 %, 2 dpl: 24,2 +/− 9,4 %, **Fig. 5D**).

**Fig. 5:**
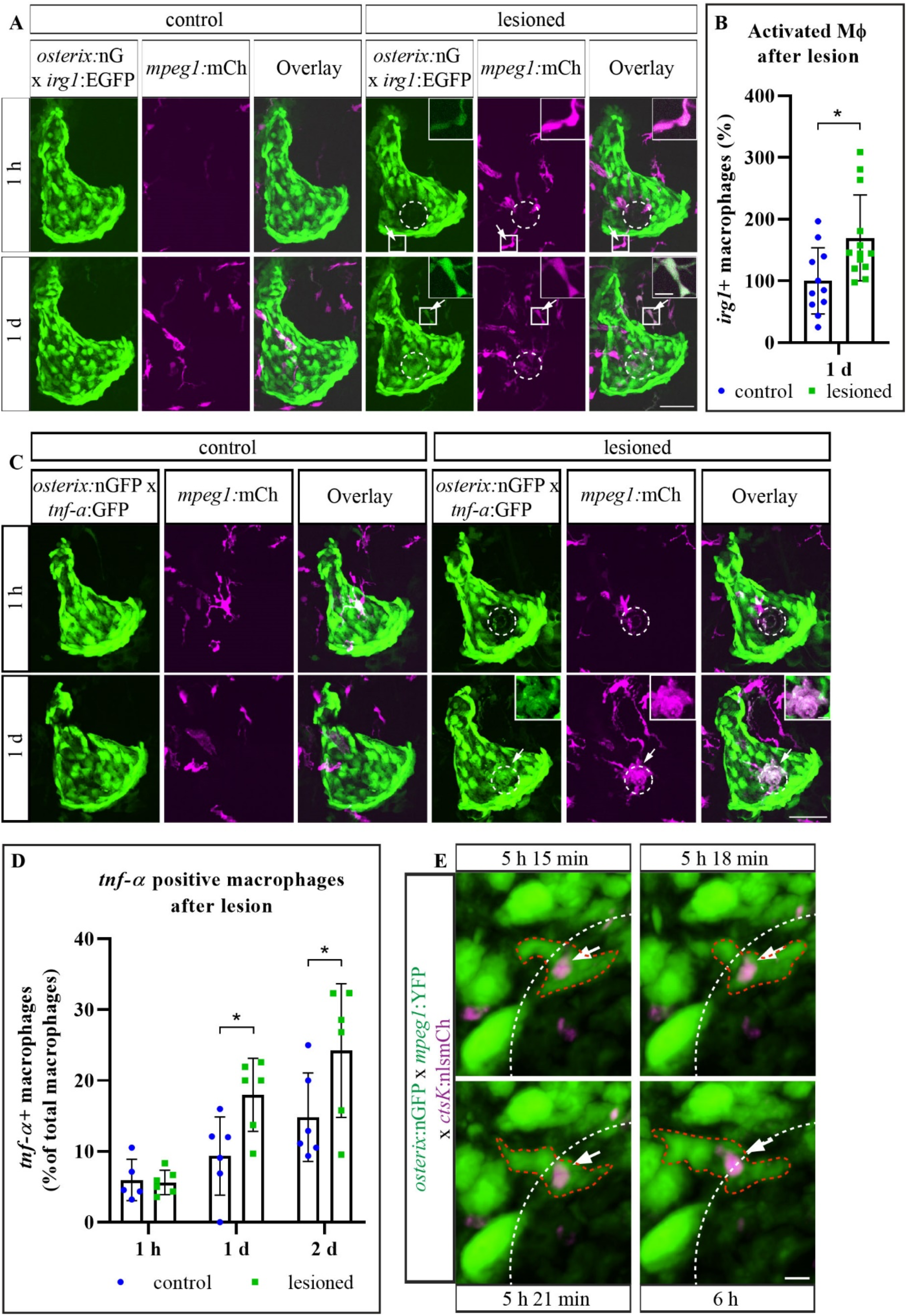
Inflammatory macrophage and osteoclast presence after osteoblast ablation. **A**: Representative images of transgenic *osterix*:nGFP x *irg*:EGFP x *mpeg1*:mCherry uninjured and ablated zebrafish opercular regions. Activated macrophages can be detected in the lesioned opercle at 1 hpl and 1 dpl by co-expression of *irg1*:EGFP and *mpeg1*:mCherry (arrows and insets). Scale bar = 20 µm. **B**: Quantification of activated macrophage numbers after osteoblast ablation in experiment shown in A. Increased numbers of activated macrophages can be detected at 1 dpl. Mean + s.d. Welch’s t-test: *p = 0.012. MФ = macrophage. n = 11-13. **C**: Representative images of transgenic *osterix*:nGFP x *tnf-α*:GFP x *mpeg1*:mCherry zebrafish opercular regions. Inflammatory macrophages can be detected in the lesioned area (white dashed line) at 1 dpl by co-expression of *tnf-α*:GFP and *mpeg1*:mCherry (arrows and insets). Scale bar = 20 µm. **D**: Quantification of inflammatory macrophages after osteoblast ablation in experiment shown in C. Increased numbers of inflammatory macrophages can be detected at 1 and 2 dpl. Mean + s.d. Sidak’s ANOVA: 1 dpl *p = 0.045, 2 dpl *p = 0.026. n = 5-6. **E**: Opercle region of transgenic *osterix*:nGFP x *mpeg1*:YFP x *ctsK*:nlsmCherry larvae at 5 to 6 hpl showing an *mpeg1*+, *ctsK*+ cell (white arrow and red dashed outline). White dashed line = border of the lesioned area. Scale bar = 5 µm. n = 4.

Next, we combined the macrophage reporter with an osteoclast reporter line established in our laboratory, in which *cathepsinK*+ cells are labeled by nuclear mCherry (*ctsK*:nlsmCherry, **Fig. S1**). This approach enabled simultaneous observation of macrophages and osteoclast-like cells after lesion. Osteoclasts are known derivatives of monocyte/macrophage lineage cells, both in mammals (Quinn et al., 1998) and medaka (Phan, Liu, et al., 2020), another teleost fish species. Using triple transgenic *osterix*:nGFP x *mpeg1*:YFP x *ctsK*:nlsmCherry zebrafish we observed YFP/nlsmCherry double positive migratory cells several hours post lesion (white arrows in **Fig. 5E**). These cells were positive for *mpeg1* and *ctsK*, indicating that macrophages convert to *ctsK*+ osteoclasts after osteoblast ablation to some extent.

These results demonstrate that ablation of approximately ten cells in a confined region is sufficient to recruit leukocytes, and that the rapid recruitment of neutrophils is followed by attraction of inflammatory macrophages. This indicates the presence of a classic early wound response in the sterile laser-assisted bone lesion paradigm. Potential conversion of macrophages into osteoclasts suggests macrophages as a source for osteoclasts in larval zebrafish.

### Antioxidant treatment suppresses macrophage attraction to the lesion site

Macrophages are attracted to their sites of action by oxidized proteins, lipids and cellular debris of apoptotic cells which are either exposed to or produce high levels of ROS (Tan et al., 2016). The presence of ROS has also been shown to be imperative for wound repair in fin fold and tail resected zebrafish larvae (LeBert et al., 2015; Romero et al., 2018). In order to assess the importance of ROS for immune cell recruitment and osteoblast recovery after laser-assisted cell ablation in bone, we treated larval zebrafish with DPI (diphenyleneiodonium chloride), a NADPH oxidase inhibitor which efficiently blocks ROS directly after fin fold amputation (**Fig. S2**) (Robertson et al., 2016), and assessed the recruitment of macrophages to the osteoblast ablation site. We observed limited recruitment of macrophages after DPI treatment **(Fig. 6, Movie 7)**, which indicates that production or release of ROS is essential for recruitment of macrophages to bone after osteoblast cell death.

**Fig. 6:**
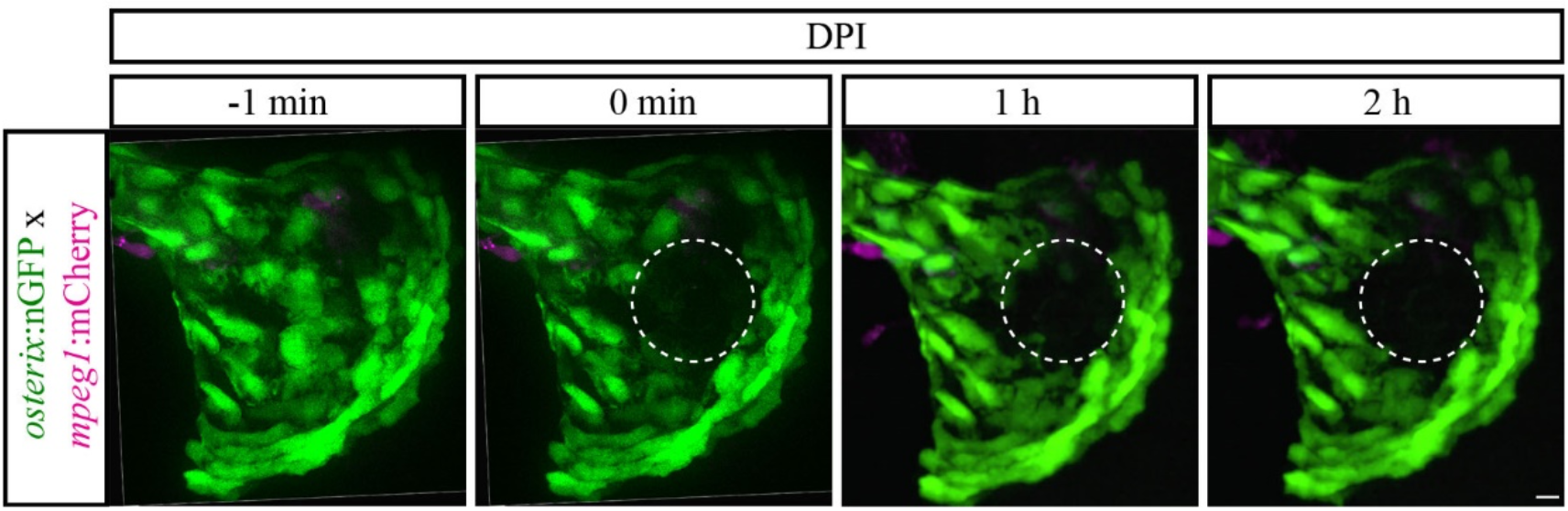
Antioxidant treatment impairs macrophage recruitment to ablated osteoblasts. Time series of osteoblast-ablated, transgenic *osterix*:nGFP zebrafish, in which macrophage recruitment into the lesioned area (white dashed line) is blocked by a 5-hour pre-treatment with the antioxidant DPI. Scale bar = 10 µm. n = 6.

### Immune-suppression by prednisolone alters macrophage recruitment to the lesion site

Macrophage activation and subtype specification need to be tightly controlled, otherwise an overexerted immune response might harm the tissue (Duffield, 2003). Persistent inflammation is also the cause for a variety of diseases that affect the skeletal system, such as rheumatoid arthritis which is routinely treated with glucocorticoids (den Uyl et al., 2011). These steroids inhibit the inflammatory response and particularly suppress macrophage recruitment in a variety of mammalian models (Cain & Cidlowski, 2017; Mosser & Edwards, 2008; Sharif et al., 2015). Making use of a previously established regime of larval zebrafish prednisolone treatment (Geurtzen et al., 2017), we tested whether mis-regulation of glucocorticoid receptor mediated signaling impacts (inflammatory) macrophage recruitment to the lesion site. After an 8-hour pre-treatment with prednisolone, which did not significantly alter the number of macrophages in the entire head of 6 dpf larvae (DMSO: 119,4 +/− 10,7 cells vs. pred: 110,4 +/− 16,92 **Fig. 7A**), and subsequent lesion, accumulation of macrophages at the lesion site was strongly reduced (**Fig. 7B, Movies 8 and 9**). A mere 10 % of the macrophages present in the opercle area were recruited into the lesion site during the first 2 hpl when prednisolone was administered, while more than 50 % of nearby macrophages were recruited to the lesion site in vehicle treated zebrafish (DMSO: 58,5 +/− 4,8 % vs. pred: 12,8 +/− 7,2 %, **Fig. 7C**).

**Fig. 7:**
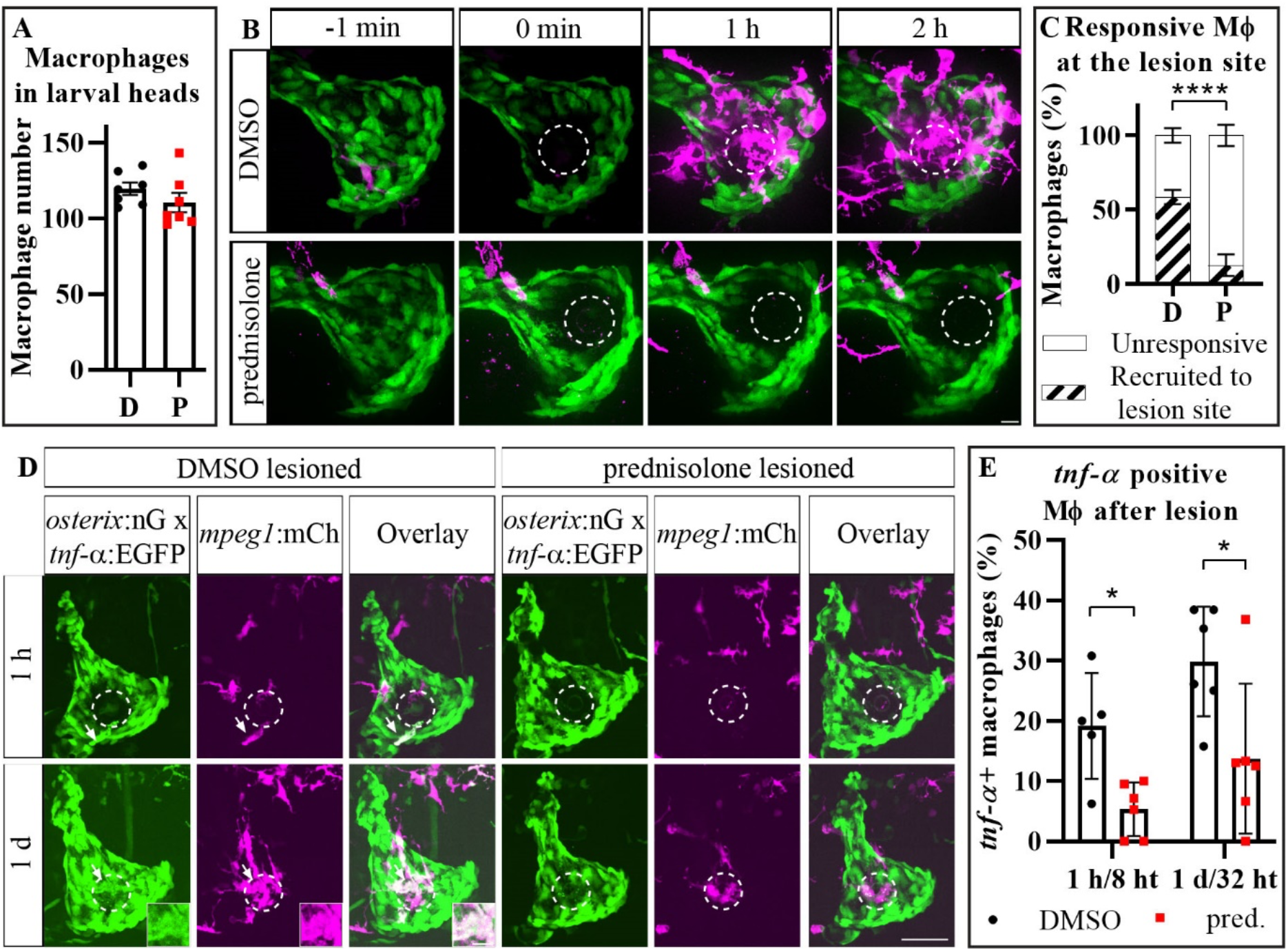
Prednisolone treatment alters the recruitment of macrophages to the ablation site. **A**: The number of macrophages in the head of 6 dpf larval zebrafish after 8 ht (hours of treatment) with prednisolone is not altered compared to the control. Mean + s.e.m. Welch’s t-test. n = 7. **B**: Time series of the opercular areas of transgenic *osterix*:nGFP x *mpeg1*:mCherry prednisolone and vehicle treated larvae. Prednisolone exposure strongly reduces the number of macrophages recruited into the lesioned area (white dashed line). Scale bar = 10 µm. **C**: Quantification of experiment shown in B. The number of responsive macrophages migrating into the lesioned area is significantly impaired by prednisolone treatment at 2 hpl. Mean + s.e.m. Sidak’s ANOVA: ****p < 0.0001. n = 6-7. **D**: Representative images of transgenic *osterix*:nGFP x *tnf-α*:GFP x *mpeg1*:mCherry osteoblast-ablated zebrafish opercular regions after prednisolone treatment. While inflammatory macrophages can be detected in the lesioned area (white dashed line) at 1 dpl by co-expression of *tnf-α*:GFP and *mpeg1*:mCherry in the DMSO control fish (arrows and insets), their presence is impaired by prednisolone treatment. Scale bar = 20 µm. **E**: Quantification of experiment shown in D. Decreased numbers of inflammatory macrophages are detected in the opercle at 1 and 2 dpl in the prednisolone treated group. Mean + s.d. Sidak’s ANOVA: 1 hpl *p = 0.044, 1 dpl *p =0.013. n = 5-6. MФ = macrophage, D = DMSO, P = prednisolone, pred. = prednisolone.

We went on to test the impact of prednisolone on the appearance of *tnf-α*:EGFP+ inflammatory macrophages and determined the respective percentage of this inflammatory macrophage subtype in the opercle after treatment. Pre-treatment with the steroid significantly reduced inflammatory macrophage numbers as early as 1 hpl (DMSO vs. pred, 1 hpl: 19,1 +/− 8,8 % vs. 5,3 +/− 4,5 %, 1 dpl: 29,8 +/− 9,1 % vs. 13,7 +/− 12,4 %, **Fig. 7D, E**). These results show that glucocorticoids severely impair macrophage recruitment to microlesions in bone tissue, simultaneously suppressing their inflammatory activated phenotype.

### Single-cell lesions allow the characterization of macrophage migratory features in response to osteoblast cell death and anti-inflammatory treatment

We asked ourselves whether smaller lesions causing cell death in fewer than 10 osteoblasts would reliably attract leukocytes to the site of lesion, a scenario potentially relevant to homeostatic tissue conditions, in which loading and cell senescence may lead to isolated cell death (Kennedy et al., 2012). In order to investigate macrophage recruitment and migration in more detail and to further study the effects of excess glucocorticoids on these features, we performed ablation of two to three osteoblasts in the center of the opercle (**Fig. 8A**, lesion outlined with white dashed line), and combined this with steroid drug administration. Recruitment of macrophages to the confined lesion site was apparent in both vehicle-treated and prednisolone-treated zebrafish (**Fig. 8B and Movies 10 and 11**); however, the relative contribution of nearby macrophages was strongly reduced compared to bigger lesions (big lesion 58,5 +/− 4,8 % vs. 18,1 +/− 8,7 % small lesion, both vehicle-treated). This indicates an injury-triggered dose-response-like mechanisms in leukocyte recruitment. In prednisolone-exposed larvae, a mild decrease of macrophage recruitment was evident (DMSO: 18,1 +/− 8,7 % vs. pred: 2,9+/− 2,9 % at 4 hpl, **Fig. 8C**), similar to what was observed after ablation of a higher number of osteoblasts. This strongly suggests that death of individual bone cells is detected by locally patrolling macrophages in otherwise unaffected tissue, and that anti-inflammatory treatment affects immune cell-osteoblast communication during tissue homeostasis.

**Fig. 8:**
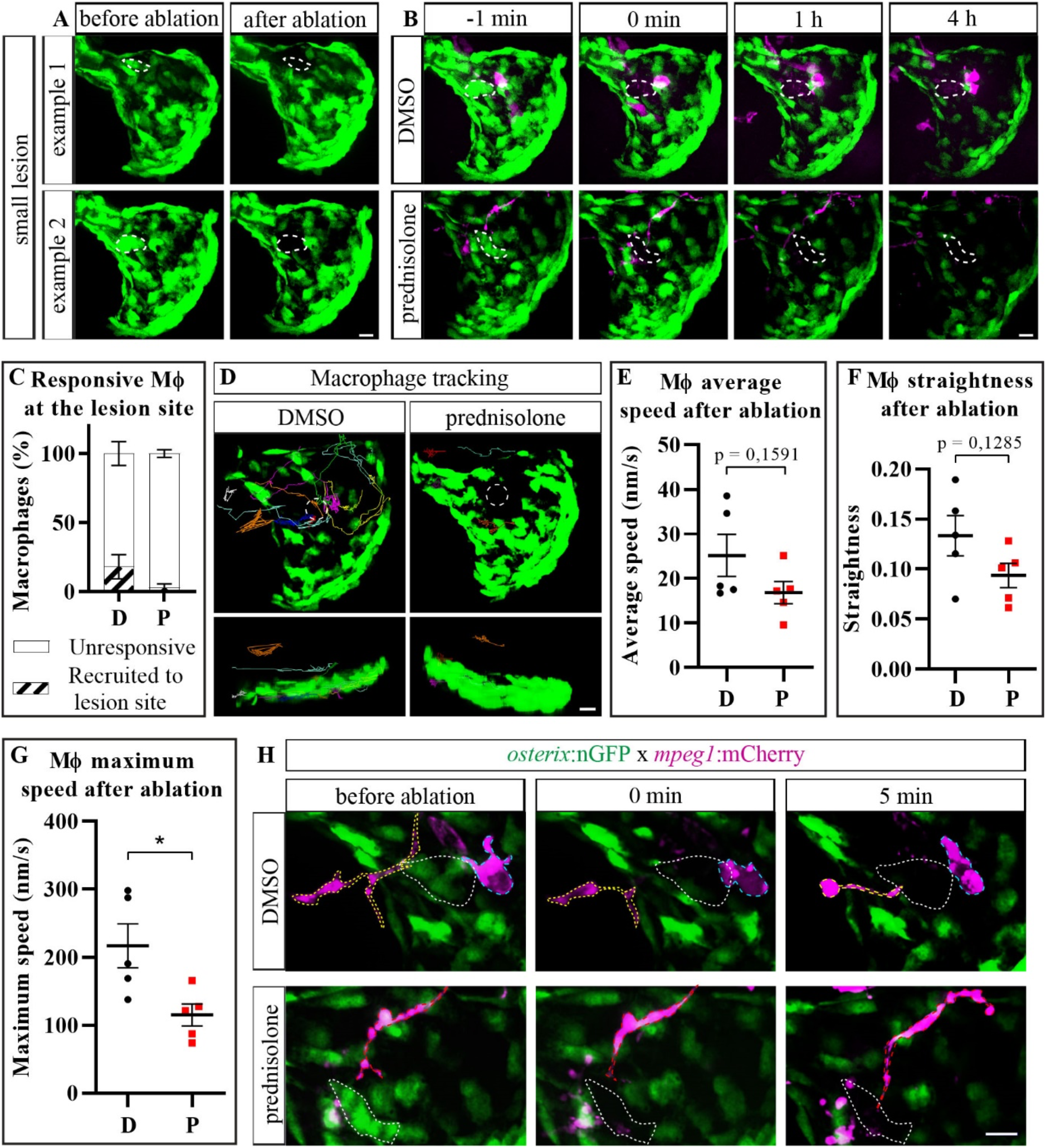
Migratory features of macrophages in response to individual osteoblast ablation and prednisolone-treatment. **A**: Examples showing specific ablation of only a few isolated osteoblast cells in vehicle treated zebrafish (precise region of osteoblast ablation = white dashed line. Scale bar = 10 µm. **B**: Time series of the opercular region in transgenic *osterix*:nGFP x *mpeg1*:mCherry vehicle treated (same as example 2 in A) vs. prednisolone treated larvae with a small lesion. Prednisolone exposure reduces the number of macrophages (magenta) recruited into the lesioned area (white dashed line). Scale bar = 10 µm. **C**: Quantification of the experiment shown in B. The response of macrophages by migration towards and into the lesion area is slightly (though not significantly) impaired by prednisolone-treatment at 4 hpl. Mean + s.e.m. Sidak’s ANOVA. n = 5. **D**: Representative images of individual macrophage tracking analysis in DMSO and prednisolone treated *osterix*:nGFP x *mpeg1*:mCherry larvae after small lesion. Arivis Vision 4D obtained tracks were overlaid with the first image after osteoblast ablation (upper panels = x-y view, lower panels = orthogonal view. Prednisolone treatment resulted in a lower number of tracks and lacking migration into the lesion site. Scale bar = 10 µm. **E**: Quantification of the average macrophage speed using the tracking shown in D. Macrophages are slightly albeit not significantly slower after prednisolone treatment. Mean + s.e.m. Welch’s t-test. n = 5. **F**: Quantification of macrophage straightness using the tracking shown in D. Straightness is slightly albeit not significantly reduced in prednisolone treated larvae. Mean + s.e.m. Welch’s t-test. n = 5. **G**: Quantification of the maximum macrophage speed using the tracking shown in D. The maximum speed of prednisolone exposed macrophages is significantly lower. Mean + s.e.m. Welch’s t-test: *p = 0.023. n = 5. **H**: Representative images of macrophages in the opercle region of *osterix*:nGFP x *mpeg1*:mCherry transgenic larvae treated with prednisolone or vehicle for 8 h. Time points shown: before, right after and 5 minutes post osteoblast ablation. White dashed line = region of ablated osteoblasts, blue dashed line = amoeboid macrophage, yellow dashed line = macrophage changing from ramified to amoeboid phenotype, red dashed line = ramified macrophage. Scale bar = 10 µm. MФ = macrophage, D = DMSO, P = prednisolone.

The lower number of recruited macrophages in microlesions enabled us to track individual macrophages by ARIVIS 4D software and to analyze migration characteristics in undisturbed, vehicle treated zebrafish larvae versus individuals after glucocorticoid treatment (**Fig. 8D**). Migratory track analysis revealed an average macrophage speed of 25,1 +/− 4,7 nm/s in vehicle treated controls, which was mildly but not significantly reduced to 16,8 +/− 2,5 nm/s by prednisolone treatment (**Fig. 8E**). Similarly, prednisolone treatment exerted subtle (albeit insignificant) effects on macrophage straightness (DMSO: 0,13 +/− 0,02 units vs. pred: 0,09 +/− 0,01 units, **Fig. 8F**), a parameter describing directional migration of cells. Importantly, glucocorticoid administered zebrafish showed significantly reduced macrophage maximum speed (**Fig. 8G**), which was decreased to about half (216,4 +/− 32,29 nm/s in DMSO vs. 115,1 +/− 16,16 nm/s in prednisolone treated individuals). This illustrates the agility of macrophages on their way to the microlesion site, and demonstrates the stationary phenotype of macrophages upon excess glucocorticoid levels.

The lower number of macrophages attracted to the lesion site also allowed a detailed investigation of macrophage morphology and respective changes upon lesion in vehicle treated versus glucocorticoid treated zebrafish. Macrophages displayed an amoeboid morphology (**Fig. 8H**, macrophage outlined in blue) or changed into an amoeboid phenotype while migrating towards the lesion site in vehicle treated individuals (**Fig. 8H**, macrophage outlined in yellow). In contrast, macrophages had a ramified and elongated phenotype with several protrusions in prednisolone-exposed individuals (**Fig. 8H**, macrophage outlined in red). These results show that ablation of individual cells triggers a considerable immune response in zebrafish bone tissue, and that short-term glucocorticoid treatment affects macrophage morphology and migration.

As prednisolone treatment impaired macrophage recruitment to the ablation site and macrophages were recently suggested to promote osteoblast differentiation and bone mineralization in mammalian bone repair (Batoon et al., 2017), we investigated the effect of prednisolone administration on recovery of osteoblast numbers after lesion. We detected a significant reduction of osteoblasts in the opercle area after lesion of prednisolone treated zebrafish (DMSO 104,7 +/− 8,43 vs. pred. 92,3 +/− 6,62 cells, **Fig. 9A**), which indicated that macrophages may have a pro-osteogenic function in the repair of microlesions. To test this, we specifically ablated macrophages by a genetic nitroreductase (NTR)-mediated killing approach (Curado et al., 2008) in triple transgenic *osterix*:CreERT2-p2a-mCherry x *hsp70*:R2nlsG x *mpeg1*:YFP-NTR zebrafish (Petrie et al., 2014), and quantified the number of lineage-traced osteoblasts in the presence or absence of NTR (schematic **Fig. 9B**). Macrophage ablated samples showed a reduced number of lineage-traced, GFP+ osteoblasts at the lesion site (NTR-2,73 +/− 0,90 vs. NTR+ 1,58 +/− 0,90 cells, **Fig. 9C, D**). In a separate experiment testing the impact of macrophage presence on general bone growth, osteoblast numbers were significantly reduced after a longer ablation period (NTR- vs. NTR+: 78,15 +/− 12,14 vs. 69,30 +/− 6,24 cells, **Fig. S3**).

**Fig. 9:**
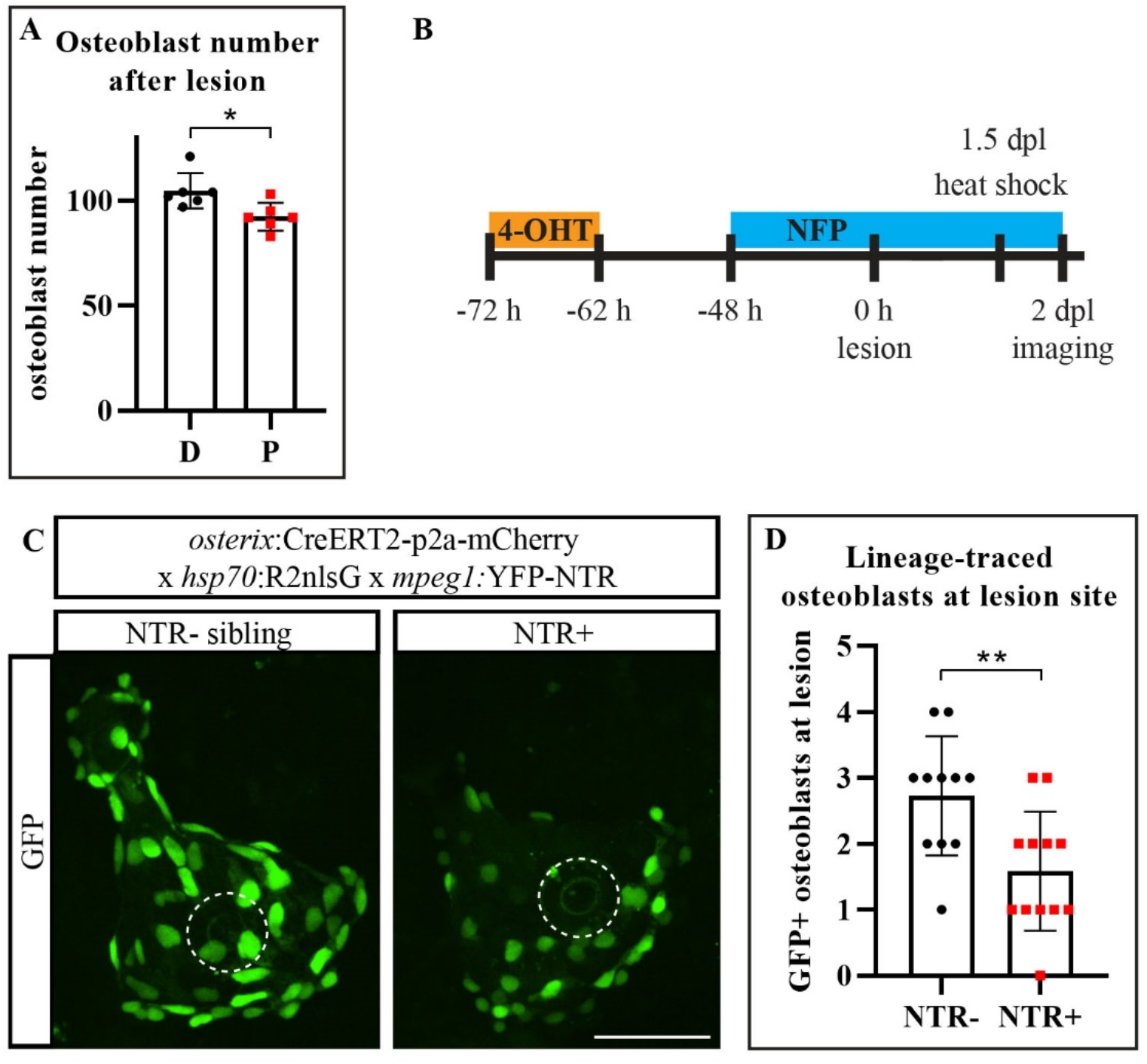
Macrophage-ablation affects the recovery of osteoblasts after ablation. **A**: Quantification of the number of opercular osteoblasts in 7 dpf transgenic *osterix*:nGFP larval zebrafish after prednisolone treatment. Mean + s.d. Welch’s t-test: *p = 0.019. n = 6. **B**: Scheme on NTR-mediated macrophage ablation combined with a CreERT2-loxP mediated lineage tracing approach of osteoblasts. Osteoblast ablation was performed at 6 dpf/72 h post 4-OHT/48 h post NFP treatment start. One day later, a single heat shock was used to visualize nuclear GFP expression. **C**: Representative images of 4-OHT and NFP-treated *osterix*:CreERT2-p2a-mCherry x *hsp70*:R2nlsG x *mpeg1*:YFP-NTR zebrafish and their NTR-siblings at 2 dpl. Macrophage ablation reduces the number of pre-existing committed opercular osteoblasts located at the lesion site (white dashed line). Scale bar = 50 µm. **D**: Quantification of experiment shown in C. Mean + s.d. Welch’s t-test: **p = 0.006. n = 11-12. D = DMSO, P = prednisolone.

In conclusion, manipulation of macrophage phenotype by pharmacologic glucocorticoid-treatment and their ablation by nitroreductase affect osteoblast recovery after microscopic bone lesion.

## Discussion

Small teleost fish such as zebrafish and medaka have proven extremely useful to monitor bone tissue during regeneration *in vivo* (Chatani et al., 2011; Cox et al., 2018; De Simone et al., 2021; Knopf et al., 2011; Phan, Liu, et al., 2020), and to observe immune cell behavior in response to infection and non-sterile soft tissue injury (Barros-Becker et al., 2017; Gray et al., 2011; Gurevich et al., 2018; Li et al., 2012). Zebrafish lesion paradigms, mostly non-sterile, have been developed for several tissues, also to study immune cell responses *in vivo* (Nguyen-Chi et al., 2015; Ohnmacht et al., 2016; Renshaw et al., 2006). Many studies make use of larval fin fold resection (’tail fin amputation’) (Demy et al., 2017; LeBert et al., 2015; Nguyen-Chi et al., 2017; Niethammer et al., 2009). The fin fold has a simple architecture, consists of two epithelial layers innervated by sensory axons, encompasses actinotrichia and interspersed mesenchyme, and lacks bone entirely (O’Brien et al., 2012). Here, we have established an approach to specifically ablate bone forming osteoblasts in larval zebrafish *in vivo*, which we used to evaluate the immune cell response towards this spatially confined, tissue-specific lesion. Our model represents a valuable tool to study the immune cell response after microscopic bone lesion and provides the possibility to evaluate the recruitment and behavior of immune cells in contributing to a balanced bone cell turnover and repair.

Recovery after cell loss is essential to ensure tissue health and the same applies to osteoblasts whose function is essential for maintenance and repair of the skeleton (X. Feng & McDonald, 2011). Many cell populations have shown the potential to generate osteoblasts, summarized under the term skeletal stem cells in mammals (Serowoky et al., 2020). In zebrafish, osteoblasts self-renew by dedifferentiation and proliferation of mature osteoblasts (Geurtzen et al., 2014; Knopf et al., 2011; Sousa et al., 2011), but also become recruited from progenitor cell pools (Ando et al., 2017; Mcdonald et al., 2021). Notably, proliferation of osteoblasts could be observed in response to laser-assisted osteoblast ablation, albeit not frequently. In addition to proliferation, stretching of cellular processes of pre-existing *osterix*+ osteoblasts towards the lesion site at 1 dpl and lineage tracing of the very same cells confirm contribution of committed osteoblasts. A rapid contractile response mediated by actomyosin forces, such as seen in the resected fin fold within less than an hour (Mateus et al., 2012), is not seen. This can potentially be explained by the comparatively firm adhesion of osteoblasts to their matrix. In the future, cell cycle studies, e.g. by labeling cells with bromodeoxyuridine and anti-phosphohistone 3 antibodies, should be performed to quantify the contribution of proliferating cells to osteoblast recovery. A potential alternative source of osteoblasts located at the lesion site may be cells that generate osteoblasts *de novo* from a precursor-like state as seen in mammals, i.e. stromal cells or other cells reminiscent of mammalian skeletal stem cells, which needs to be tested in the future.

We did not detect developmental delays of bone formation after osteoblast lesion. This might be because of quick osteoblast recovery and due to the fact that opercular growth at the investigated developmental stages is comparatively slow; the volume of the opercular bone matrix increases only by about 10 % from 7 to 8 dpf (**Fig. 1C**). Furthermore, opercular growth mostly proceeds along the ventral-posterior bone edge with osteoblasts strongly concentrating in this region (Kimmel et al., 2010) which was also enriched for lineage-traced osteoblasts (**Fig. 2C**). The ventral-posterior edge of the opercle was not targeted in our lesion paradigm. It will be interesting to test whether osteoblast ablation in this region leads to a growth defect, and if so, how many osteoblasts would need to be ablated to evoke alterations.

One important process after wounding is the production and release of cytokines attracting immune cells, which is essential to initiate tissue repair (Duffield, 2003). Mmp9, one of the signals induced after tissue damage (LeBert et al., 2015; Matsubara et al., 1991), triggers leukocyte migration (Purwar et al., 2008). Similarly, high ROS amounts are released after wounding (Roy et al., 2006; Yoo et al., 2012) and stimulate recruitment of immune cells in zebrafish (Y. Feng et al., 2010; Niethammer et al., 2009). Sustained ROS levels are also observed after adult fin amputation, which includes the resection of bone (Gauron et al., 2013), and after resection of the larval notochord (Romero et al., 2018). Both *mmp9* activity and ROS production were induced by sterile laser-mediated osteoblast ablation as established in this work, which illustrates that ablation of less than a dozen cells causes a wound response comparable to the one seen after non-sterile, more severe wounding. However, *mmp9* activity was visualized with the help of a transgenic reporter, which requires several hours to mirror transcript activity of the *mmp9* gene, during which GFP protein is produced (Hazelrigg et al., 1998). For this reason, we cannot infer how fast *mmp9* message and protein are formed and whether Mmp9 release indeed plays a role in early leukocyte attraction. Conversely, rapid ROS release was visualized real time, and treatment with the antioxidant DPI reduced leukocyte attraction, indicating that ROS are involved. Classic work on zebrafish fin fold resection (Niethammer et al., 2009) and tumor-transformed skin cells (Y. Feng et al., 2010) demonstrated the importance of ROS-mediated immune cell attraction, which is likely to occur short-range (Jelcic et al., 2017). ROS blockage in our experiments may either cause reduced ROS levels at the ablation site or reduce ROS levels in macrophages, and therefore lead to impaired recruitment. Notably, ROS accumulate in macrophages and other leukocytes playing an important role in leukocyte polarization (Robinson, 2008; Tan et al., 2016). Moreover, ROS production plays a crucial role in bone homeostasis promoting osteoclastogenesis and bone resorption (Bai et al., 2005; Lee et al., 2005), and oxidative stress is strongly associated with bone pathologies such as osteoporosis (Baek et al., 2010). However, to our knowledge ROS have not been visualized at cellular resolution *in vivo* in bone before, making this the first study to successfully label ROS release after osteoblast cell death in a living organism. It remains unclear, whether increase of ROS in or by individual osteoblasts is sufficient to attract immune cells. In the future, new tools such as KillerRed will enable tissue and cell type specific generation of ROS (Formella et al., 2018; Teh et al., 2010) in the growing opercle without ablating the cells. This will lead to the further characterization of the impact of ROS on bone cell turnover and tissue homeostasis.

In case of tissue damage a rapid immune cell influx is essential for efficient cell and tissue clearance and to ensure proper inflammation resolution and healing (Duffield, 2003). Both neutrophils and macrophages were recruited to the site of laser-assisted osteoblast ablation. Neutrophils were recruited first, and macrophages followed shortly after, which is in agreement with previous work (Keightley et al., 2014; Kolaczkowska & Kubes, 2013). The observed macrophages were tissue-resident, responded early, and might be responsible for later recruitment of monocyte-derived macrophages and other inflammatory leukocytes to the injury site (Davies et al., 2013). Moreover, the average speed of macrophages was comparable to previously reported zebrafish injury models (Barros-Becker et al., 2017; Ellett et al., 2011; Li et al., 2012). Furthermore, acquisition of *ctsK* expression in some macrophages, indicating osteoclast differentiation, was observed. A similar process was suggested to take place in medaka fish, in which RANKL (Receptor activator of NF-κB ligand) was ubiquitously overexpressed (Phan, Liu, et al., 2020; Phan, Tan, et al., 2020). Some caution is warranted, as *ctsK* also labels mesenchymal cells in some tissues (Debnath et al., 2018; Lu et al., 2020) and we observed immobile *ctsk*:nlsmCherry+ nuclei at a distance from the opercle, in locations unlikely to contain osteoclasts (data not shown). It is noteworthy that conversion of macrophages to osteoclasts depends on inflammatory signals such as Tnf-α (Phan, Liu, et al., 2020). We detected increased expression of *tnf-α* in some macrophages recruited to the lesion site, and these cells are good candidates for differentiation into osteoclasts. Further investigation of additional osteoclast-specific and inflammatory marker gene expression, such as of tartrate-resistant acid phosphatase, will reveal the importance of macrophage recruitment and inflammatory phenotype for osteoclastogenesis at the lesion site.

Macrophage contribution to wound healing is dependent on the subtype characteristics. Generally, the early wound-response is characterized by inflammatory macrophage action marked by inflammatory cytokine release and phagocytosis of dead cells and debris (Duffield, 2003; Lieschke et al., 2001). Later phases are dominated by regenerative macrophage responses leading to inflammation resolution and tissue remodeling (Duffield, 2003; Novak & Koh, 2013). In zebrafish, transgenic reporter lines enable investigation of macrophage phenotypes throughout inflammation and its resolution (Ellett et al., 2011; Gray et al., 2011; Walton et al., 2015). Available tools showed that, similarly to the mammalian situation, two waves of macrophages (inflammatory and regenerative) emerge in soft zebrafish tissues after resection (Nguyen-Chi et al., 2017).

In this work, we evaluated the amount of activated and inflammatory macrophages at relatively early time points post cell ablation. Interestingly, increased *irg1*:GFP+ macrophages were already detected at 1 hpl indicating that these activated macrophages respond fast. While Irg1 is not a direct read-out for the inflammatory macrophage phenotype, but instead labels activated macrophages (Sanderson et al., 2015), it is often expressed in a pro-inflammatory environment (Jamal Uddin et al., 2016) by pro-inflammatory macrophages (Sanderson et al., 2015). We therefore conclude that activated macrophages populate the lesion site early on and that they become polarized towards the inflammatory phenotype. Notably, only about 25 % of macrophages in the opercle area and at the lesion site were identified as inflammatory macrophages at 1 dpl, on the basis of *tnf-a*:GFP expression. This is in agreement with work on fin fold resected zebrafish, in which approximately 30 % of inflammatory macrophages were reported at 24 hours post amputation, although more inflammatory macrophages could be detected earlier (Nguyen-Chi et al., 2017). In our model, laser-assisted osteoblast ablation will allow to study the kinetics of the inflammatory response to cell death in bone tissue, in particular with respect to normal and dysregulated resolution of inflammation.

Tight regulation of the immune response is essential for wound healing and tissue repair, and overexerted actions can be harmful in many inflammatory disorders (Duffield, 2003). In the bone microenvironment, a variety of inflammatory conditions and therapy-associated diseases such as rheumatoid arthritis and glucocorticoid-induced osteoporosis affect bone health (den Uyl et al., 2011; X. Feng & McDonald, 2011; Takayanagi, 2007). In order to develop novel therapies for such diseases and to counteract adverse effects of immunosuppressive treatment, it is crucial to precisely understand how immune cells are affected in these conditions. Zebrafish fin fold regeneration models combined with high dose glucocorticoid treatment (Chatzopoulou et al., 2016; C. J. Hall et al., 2014; Sharif et al., 2015) showed that glucocorticoids suppress the attraction of macrophages and neutrophils and impair tissue regeneration. Here, by using a previously established larval prednisolone administration regime (Geurtzen et al., 2017) on laser-ablated zebrafish larvae, we investigated the effect of glucocorticoids on immune cells *in vivo* in the context of bone cell turnover and microscopic bone repair. Particularly inflammatory macrophages were affected by prednisolone exposure, as reported previously (Cain & Cidlowski, 2017; Russo-Marie, 1992; Xie et al., 2019). Morphology, which can be used as a readout for activation and polarization status of macrophages (McWhorter et al., 2013), was evidently altered after prednisolone exposure. Inflammatory, activated macrophages display an amoeboid morphology with few dendrites while anti-inflammatory macrophages are more elongated and display more dendrites, also in zebrafish (Nguyen-Chi et al., 2015). Prednisolone exposure impaired the activated, amoeboid morphology of macrophages in our model. Consistently, *tnf-α*:GFP+ macrophage numbers dropped in prednisolone treated zebrafish, similar to effects after fin fold resection and concomitant treatment (Nguyen-Chi et al., 2015).

Contrasting data have been obtained concerning the effect of glucocorticoids on immune cell migration, also varying depending on the glucocorticoid used (Chatzopoulou et al., 2016; Sharif et al., 2015; Xie et al., 2019). While beclomethasone leaves macrophage migration unaffected in fin fold resected zebrafish (Chatzopoulou et al., 2016; Xie et al., 2019), high dose dexamethasone treatment reduces macrophage recruitment in the same model (Sharif et al., 2015). Here, we used prednisolone, a third synthetic glucocorticoid, to test for its immunosuppressive effects after individual osteoblast ablation. Treatment led to a reduced macrophage migratory ability in terms of maximum speed and number of recruited macrophages. This is in agreement with previous results on adult bony fin ray-amputated zebrafish, in which prednisolone treatment led to impaired macrophage accumulation in fin regenerates (Geurtzen et al., 2017).

High dose glucocorticoids are known to strongly affect osteoblasts by inducing osteoblast apoptosis while impairing osteoblast proliferation and maturation (den Uyl et al., 2011; Weinstein, 2012). To date, it is unclear whether these anti-osteogenic effects are exclusively mediated directly or whether alteration of macrophage number and phenotype contribute to these effects. During mammalian bone repair, macrophages promote osteoblast differentiation and bone mineralization (Pettit et al., 2008), in particular after fracture (Batoon et al., 2017). In our lesion model, ablation of macrophages led to reduced osteoblast numbers at the lesion site, reflecting either impaired proliferation or migration of *osterix*+ osteoblasts. Decreased osteoblast numbers in the developing opercle after macrophage ablation point to a similar compromising effect, which was also observed in fractured mouse bones after tissue-resident macrophage ablation (Alexander et al., 2011; Batoon et al., 2017). This illustrates the capacity of tissue-resident macrophages to support bone formation in mammalian and non-mammalian vertebrates. In the future, it will be interesting to test osteoblast and macrophage-specific knockout tools in zebrafish, e.g. to delete the glucocorticoid receptor *nr3c1*, or to target prednisolone to phagocytic macrophages specifically to decipher the indirect negative impact of glucocorticoid exposure on osteoblasts via immune cells. Moreover, it will be interesting to compare anti-migratory effects of different synthetic glucocorticoids on macrophages and neutrophils across different wounding assays and tissues, specifically when taking bone tissue into account.

*In vivo* and intravital imaging approaches in rodent species have progressed a lot in recent years. Tissues such as the lung (Yang et al., 2018), kidney (Peti-Peterdi et al., 2016) but also bone (J. Kim & Bixel, 2020) have been investigated. A variety of studies examined the interaction of bone producing cells with immune cells by using *in vivo* microscopy (Tetsuo Hasegawa et al., 2019; Ishii et al., 2010; Kikuta et al., 2013). Ishii and colleagues used two-photon confocal laser microscopy to observe osteoclast precursor migration to bone tissues in homeostatic conditions (Ishii et al., 2009). Similar approaches, some of which making use of bone explants, facilitated imaging of osteoblast – osteoclast interactions, osteoprogenitors during cranial bone defect repair and the mechanism of osteocyte embedding into bone ECM (Dallas & Moore, 2020; Furuya et al., 2018; Huang et al., 2015; Shiflett et al., 2019). Albeit these advancements, limiting factors in terms of imaging depth persist for *in vivo* imaging of rodent bone tissue. Furthermore, the ability to resolve cellular dynamics in terms of cell shape changes, migratory behavior and cell to cell contacts remains challenging. This is also true for long-term imaging of rodent bone tissue *in vivo*, which, in contrast, can be performed up to several days in zebrafish larvae (Kaufmann et al., 2012).

Laser-assisted approaches have a long tradition in zebrafish since they provide good tissue penetration (Morsch et al., 2017) and fine spatio-temporal control of manipulation (Johnson et al., 2011). This makes the system useful for a wide array of experiments triggering regeneration, such as by laser-induced axotomy (Hu et al., 2018), induction of thrombosis (Jagadeeswaran et al., 2006), cell ablation in the brain (Sieger et al., 2012), spinal cord (Dehnisch Ellström et al., 2019), kidney (Johnson et al., 2011), intestine (Ohno et al., 2021) and heart (Matrone et al., 2013). In bone, only few laser-assisted ablation assays have been employed. Two assays were performed in non-osseous tissues - the wound epidermis covering regenerating bone elements in fins (J. Zhang et al., 2012) and hypothalamic neurons during early development (Suarez-Bregua et al., 2017), which lead to altered fin ray patterning and mineralization defects in craniofacial bones, respectively. Substantial osteoblast ablation was performed by Chang & Franz-Odendaal in selected skeletal condensations of developing infraorbital bones in older zebrafish larvae (10 mm standard length, age > one month, (Singleman & Holtzman, 2014)). While the bones recovered, they were however smaller and aberrant in shape (Chang & Franz-Odendaal, 2014). In our assay, ablation of a maximum of 10 % of opercular osteoblasts was subcritical and did not compromise bone growth, which could also reflect the ability of younger individuals to better compensate for tissue damage during development.

Our model represents a novel approach to study bone cell turnover in homeostasis and repair. We show that ablation of only few osteoblasts is sufficient to initiate an immune cell response of graded severity, depending on how many cells were ablated initially. This is especially interesting considering that osteoblast senescence and cell death are frequent processes in bone homeostasis with osteoblasts having a relatively short life span (Manolagas, 2000). The consequences of isolated osteoblast death during homeostasis and medical treatment, as well as their replacement, are not particularly clear. In recent years, osteocytes have gained center stage as sensors of biomechanical strain and players in modeling and remodeling of bone (BONEWALD, 2007; Kennedy et al., 2012; Ru & Wang, 2020). However, osteoblasts, mirroring the pre-osteocytic cell stage, have suggested to play a similar role in sensation of microscopic tissue damage. This is illustrated by the fact that anosteocytic bones undergo modeling in teleost fish (medaka (Ofer et al., 2019)), which is mediated by Sclerostin levels in bone lining osteoblasts, and is supported by the observation that osteocytic bones do not necessarily undergo remodeling (Currey et al., 2017). Another aspect of osteoblast biology is their declined performance in aged bone, which is partly mediated by accumulation of ROS (H. N. Kim et al., 2018), and rising levels of glucocorticoids (M Almeida & O’Brien, 2013). Both factors, which we modeled with the help of laser-assisted osteoblast ablation, influence bone health directly and indirectly, by impairing osteo-immune cell communication (Ahmad et al., 2019; Lean et al., 2005). While we are aware that the presented assay makes use of a growing bone of simple structure, we suggest to use it to *in vivo* monitor the processes of osteoblast recovery after cell death, and to study the varying contribution of immune cells to bone repair and integrity. In addition, interaction of leukocyte cell types, such as the process of reverse migration of neutrophils first observed in zebrafish (Mathias et al., 2006) and later demonstrated in mammals (Woodfin et al., 2011), and the generation of macrophage/monocyte-derived osteoclasts in response to ROS and increased stress signaling can be studied. Finally, the model enables studies on the contribution of different osteoblast progenitor cell pools to osteoblast recovery and can potentially be used in slightly older animals to visualize the plasticity of mature osteoblasts undergoing dedifferentiation.

## Summary

In conclusion, we present a comprehensive study on a new laser-assisted osteoblast ablation paradigm, which enables researchers to study osteoblast – immune cell interactions *in vivo*. We investigated the specific characteristics of this sterile lesion assay and show that a varying number of osteoblasts can be ablated and that recovery occurs fast when 10 % of the opercular osteoblasts are ablated. Using spinning disc confocal microscopy, we tracked the immediate response of neutrophils and macrophages, which migrated into the site of ablation at which ROS were released quickly. A significant number of recruited macrophages displayed an inflammatory phenotype, which was inhibited by pharmacological glucocorticoid exposure. Moreover, glucocorticoid-treatment significantly impaired macrophage migration into the region of interest and affected osteoblast recovery. Ablation of *mpeg1*+ macrophages impaired osteoblast repopulation of the injured area suggesting a bone-anabolic macrophage function. Laser-assisted ablation of osteoblasts can be used to better understand microscopic bone repair during tissue homeostasis and to explore the relevance of leukocyte recruitment in this process.

## Material & Methods

### Animal experiments

All procedures were performed in accordance with the animal handling and research regulations of the Landesdirektion Sachsen (Permit numbers AZ DD25-5131/354/87, DD25-5131/450/4, 25-5131/496/56 and amendments).

### Fish lines and husbandry

The following previously described transgenic zebrafish lines were used: *osterix*:nGFP (Tg(*Ola.Sp7*:NLS-GFP)^zf132^)(Spoorendonk et al., 2008), histone Cherry (Tg(*h2afv*:h2afv-mCherry)^tud7^)(Knopf et al., 2011), *mmp9:*EGFP (TgBAC(*mmp9*:EGFP-NTR)^tyt206^)(Ando et al., 2017), mpo:GFP (Tg(BAC*mpo*:gfp)^i114^)(Renshaw et al., 2006), m*peg1:*mCherry (Tg(*mpeg1*:mCherry)^gl23^)(Ellett et al., 2011), *mpeg1:*YFP-NTR (Tg(*mpeg1*:NTR-EYFP)^w202^)(Petrie et al., 2014), TgBAC(*tnfa*:GFP)^pd1028^)(Marjoram et al., 2015), *irg1:*EGFP (Tg(*acod1*:EGFP)^nz26^)(Sanderson et al., 2015), *osterix*:CreERT2-p2a-mCherry (Tg(*Ola.sp7*:CreERT2-P2A-mCherry)^tud8^)(Knopf et al., 2011), *hsp70*:R2nlsG (Tg(*hsp70l*:loxP-DsRed2-loxP-nlsEGFP)^tud9^)(Knopf et al., 2011), *kdrl*:CFP (Tg(*kdrl*:CFP)^zf410^)(Hess & Boehm, 2012).

For creation of the *ctsK*:nlsmCherry transgenic zebrafish line a fragment containing the 4 kb promotor region and start of exon1 of the *Danio rerio cathepsin K* was cloned upstream of nlsmCherry into a pBluescript-based vector containing Tol2 transposable sites flanking the insert (kindly provided by Anke Weber and Stefan Hans). The following primers were used for amplification: ATATCCTCTCACAGGACATCAAACAGCGAAACGAG (adding an EcoNI restriction site) and TATAGGCCGGCCTGAGCAAGAAGAAATGCACC (adding a FseI restriction site). EcoNI and FseI restriction enzymes were used for cloning. A transgenic line was created by injecting the plasmid DNA with transposase mRNA into fertilized eggs. Throughout larval growth transgene expression was detectable in the pharyngeal region comparable to another previously published zebrafish *ctsK*-line (Sharif et al., 2014).

Fish were bred and maintained as described (Brand M et al., 2002).

### Lesion paradigm and *in vivo* imaging

In order to perform osteoblast ablations, zebrafish larvae were anesthetized in 0.02 % Tricaine (MS222, Merck, Taufkirchen, Germany) and embedded in 1 % Low melt agarose (LMA, Biozym Scientific GmbH, Hessisch Oldendorf, Germany) in E3 in a glass bottom microwell dish (35 mm, 14 mm microwell, MatTek Corporation, Ashland, MA, USA). To immobilize the larvae, 20 µl of 0,4 % Tricaine were added to 1.5 ml LMA (final concentration 0.005 % Tricaine). They were laid on the side to position the opercle region close to the glass bottom for imaging accessibility. For prednisolone-treatment the larvae were kept in prednisolone (Merck, Taufkirchen, Germany) or DMSO (Merck, Taufkirchen, Germany) control conditions by adding autoclaved E3 with 0.01% Tricaine and 25 µM prednisolone or 0.05% DMSO to the dish after the LMA had solidified, otherwise E3 with 0.01% Tricaine was added. The osteoblast lesion was performed using a UV laser adjusted to the Andor spinning disk system equipped with a Yokogawa CSU-X1, Zeiss AxioObserver.Z1 and an iXon+ camera facilitating simultaneous imaging with LabVIEW 2009. The same laser cutter settings were used throughout the study, consisting of 2.0 intensity, 20 pulses/shot, four shots/µm^2^ and two shooting circles of 15 and 7 µm Ø. Cell death was confirmed by loss of GFP signal. Afterwards the larvae were carefully removed from the LMA and kept in E3 until further imaging was performed. Non-lesioned controls were mock-treated i.e. they were also anesthetized and embedded in LMA.

Short-term imaging was performed with LabVIEW 2009 with the setup described above, while long term imaging was performed with Andor iQ2 software or a Dragonfly spinning disk equipped with a sCMOS camera and Fusion software. In both imaging approaches, the same laser power, gain settings and exposure times were used. Larvae were kept in LMA after the lesion and were continuously imaged at different time intervals. Data were processed with Image J Software version 1.53c or Arivis Vision4D version 2.12.6. and 3.3.0 where mentioned, in order to obtain images and movies. For tracking of macrophages Arivis Vision4D version 2.12.6. was used. First, the data were filtered using a convolution enhancement filter which was followed by drift correction using the GFP channel. Afterwards, individual macrophages were tracked using the blob finder and Brownian motion segment tracker.

Only macrophages whose complete cell bodies were visible and which very present in the field of view for more than 30 min were tracked. At the end, the tracking was manually verified, aberrant tracks were excluded and separated tracks were merged.

### Drug treatments

Prednisolone treatment of zebrafish larvae was carried out as previously described (Geurtzen et al., 2017).

DPI (Diphenyleneiodonium chloride, Merck, Taufkirchen, Germany) treatment of 3 dpf zebrafish larvae before finfold resection was carried out as previously described (Robertson et al., 2016). DPI treatment on 6 dpf larvae started 5 h before laser ablation and continued throughout the live imaging period with the same concentration used on 3 dpf larvae (100 µM).

Nifupirinol (NFP) treatment was performed as previsouly described (Bergemann et al., 2018). A 2.5 mM stock solution of NFP in DMSO was prepared and stored at −20°C. Larval zebrafish were soaked in 2.5 μM NFP for up to 6 consecutive days in the dark at 28°C.

### Fin fold resection

3 dpf larvae were anesthetized and fin fold resection was performed as previously described (Isles et al., 2019).

### Staining techniques

Alizarin red staining of the live zebrafish larvae was performed as previously described (Kimmel et al., 2010). For quantification of the opercle volume the surface tool in Imaris 8.1 was used to reproduce the alizarin red stained area surface and to calculate the corresponding volume.

CellROX staining was conducted as previously described (Kulkarni et al., 2018), however, instead of CellROX green, CellROX orange (ThermoFisher Scientific, Waltham, MA, USA) was used. After staining, larvae were embedded in 1% LMA, lesioned and live imaged for 15 min.

### Osteoblast fate mapping

*osterix*:CreERT2-p2a-mCherry, *hsp70*:R2nlsG double transgenic 5 dpf larvae were soaked in 10 µM 4-hydroxytamoxifen (4-OHT, Merck, Taufkirchen, Germany) or the corresponding amount of vehicle control ethanol for 10 h. Larvae were lesioned at 6 dpf. One or two days later (7 or 8 dpf), larvae were heat shocked once at 37°C for 1 h. Roughly 12 h post heat shock the larvae were analyzed for recombination by the appearance of nlsGFP+ cells.

### Quantification of GFP and mCherry expression, steromicroscopy

For quantification of GFP and mCherry expression post lesion the larvae were anesthetized with 0.02% Tricaine (MS222) and again embedded in 1% LMA in a glass bottom dish. The larvae were imaged once with an Andor Dragonfly Spinning Disk equipped with a sCMOS camera. For each time point a different set off larvae was used. Identical settings for magnification, exposure time, pinhole size, laser power and z-stack interval were used throughout the whole experiment. Intensity measurements were conducted using the Plot Profile Tool in Image J Software version 1.53c.

Stereomicroscopy of fin fold resected zebrafish larvae and the head region of *ctsK*:nlsmCherry transgenic zebrafish was performed with the help of a Zeiss SteREO Discovery.V12 equipped with a AxioCam MRm and AxioVison software version 4.7.1.0.

### Image processing

Brightness, contrast and levels were adjusted using Adobe Photoshop CS6 and 2020 software. Images were processed with identical settings using the legacy option.

### Statistical analysis

Statistical analysis was run using GraphPad Prism 8.3.1. Unpaired two-sided t-tests with Welch’s correction, Tukey’s multiple comparison one-way ANOVA and Sidak’s multiple comparison two-way ANOVA tests were performed wherever applicable.

## Acknowledgements

We would like to thank Isabell Weber for help in establishing the laser-assisted lesion assay and Michael Brand for helpful discussions. We are very grateful to Atsushi Kawakami (Tokyo, Japan), Michel Bagnat (Durham, UK) and C.J. Hall (Auckland, New Zealand) for sharing transgenic fish. Special thanks goes to Stefan Grill, Nicholas Chartier and Lokesh Pimpale for providing access to their Spinning Disk Microscope and technical support. We would also like to thank Anke Weber and Stefan Hans for sharing reagents. Our thanks also goes to the Light Microscopy core facility at the Center for Molecular and Cellular Bioengineering at the TU Dresden and Marika Fischer, Jitka Michling and Daniela Moegel for excellent fish care. We thank Henriette Knopf for proofreading the manuscript.

## Author contributions

Experiments were designed and analyzed by KG, AD and FK. KG and AD performed experiments. KG and FK wrote the manuscript and accept responsibility for the integrity of data analysis.

## Supplemental Information

**Fig. S1:**
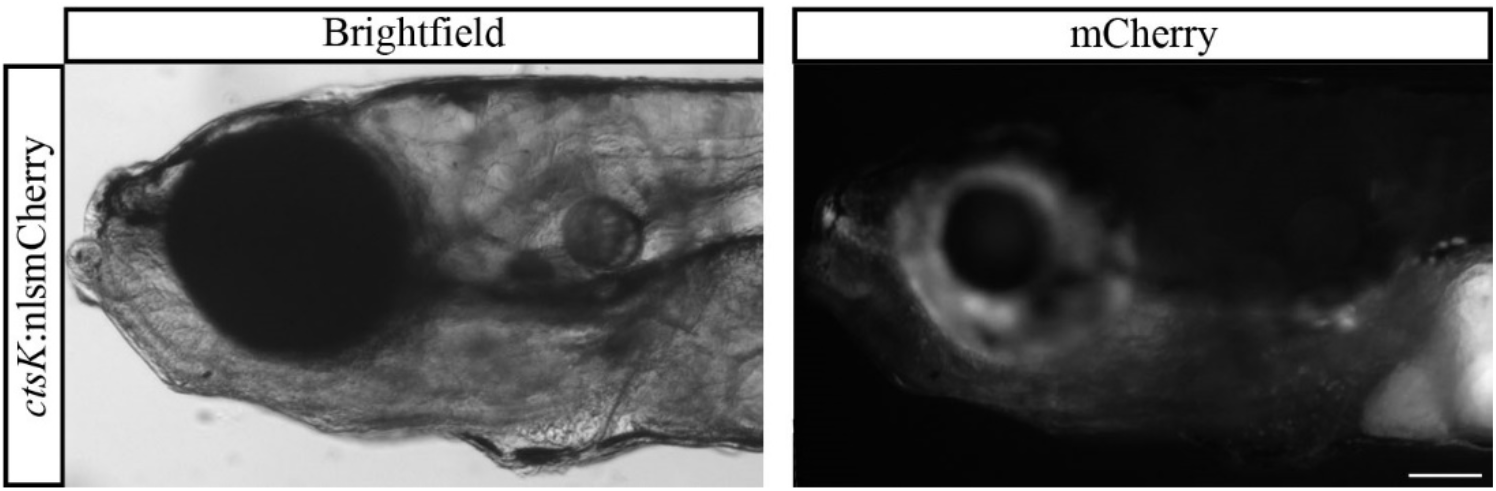
Representative whole-mount image of transgenic *ctsK*:nlsmCherry larval heads at 13 dpf. The staining is visible in the whole lower jaw region. Scale bar = 100 µm. n = 2

**Fig. S2:**
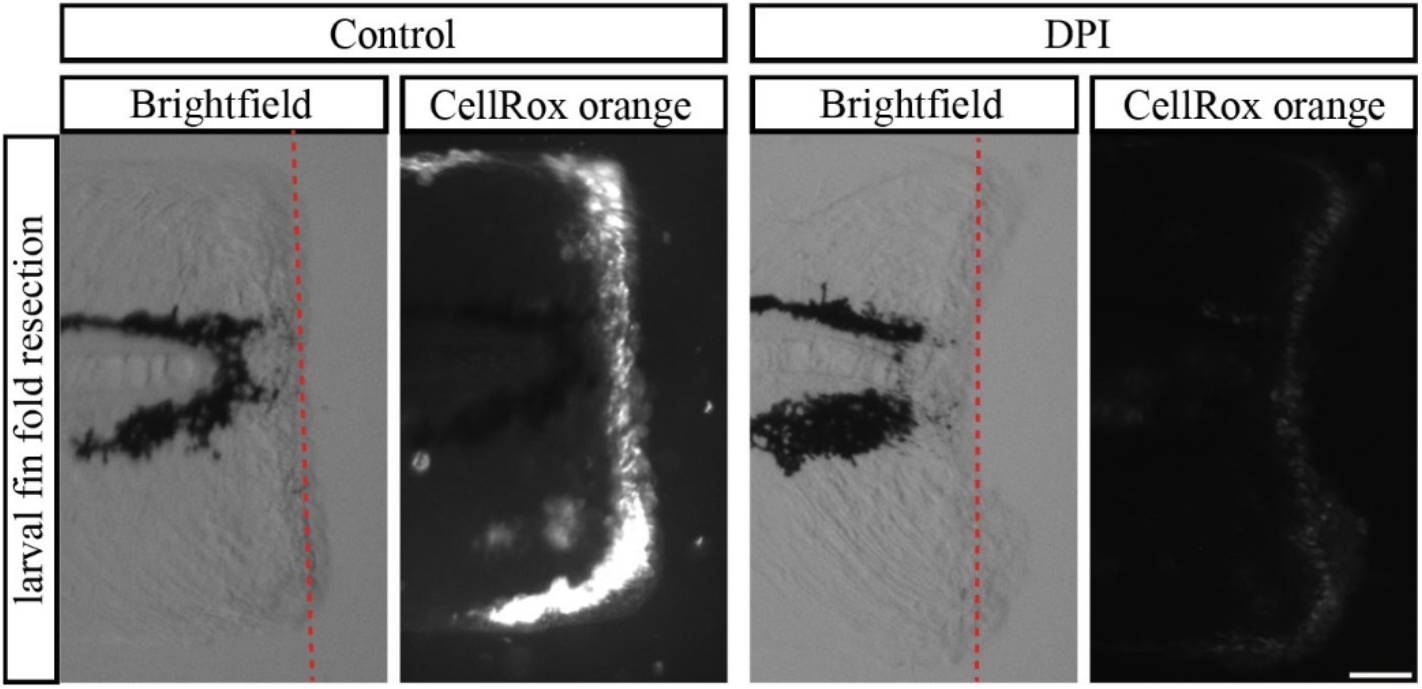
Representative whole-mount images of ROS production, indicated by CellROX orange staining, 20 min after fin fold resection. The release of ROS can be blocked by a pre-treatment with the antioxidant DPI (diphenyleneiodonium). Scale bar = 50 µm. n = 6

**Fig. S3:**
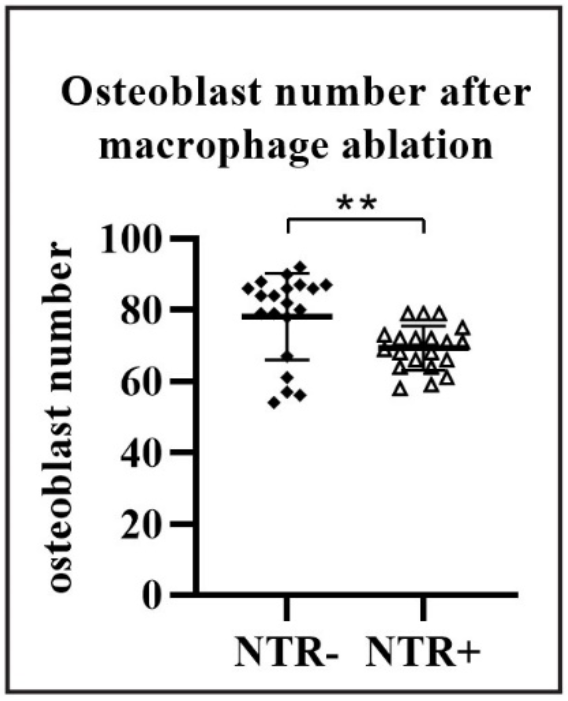
Quantification of the number of osteoblasts on the uninjured opercle after ablation of macrophages with NFP. *osterix:*RFP x *mpeg1:*YFP-NTR larvae and *mpeg1*:YFP-NTR negative siblings were incubated for 6 days with NFP. A reduced number of osteoblasts at 9 dpf was detected in case of macrophages ablation. n = 20.

**Movie 1 (supplement to Fig. 2):** Movie showing a proliferating osteoblast in the opercle *of osterix*:nGFP transgenic zebrafish after osteoblast ablation. The position of the proliferating osteoblast is indicated by the white arrow. Division can be observed after roughly 4 h. Scale bar = 10 µm.

**Movie 2 (supplement to Fig. 3C):** Movie showing absence of CellRox orange staining in a transgenic 6 dpf *osterix*:nGFP zebrafish with osteoblast ablation. Scale bar = 10 µm.

**Movie 3 (supplement to Fig. 3C):** Movie showing increasing CellRox orange staining, as a readout for ROS release, in a transgenic 6 dpf *osterix*:nGFP zebrafish after osteoblast ablation. The increase of ROS is observed immediately lesion. Scale bar = 10 µm.

**Movie 4 (supplement to Fig. 4A):** Movie showing the recruitment of neutrophils, labeled in magenta, into the area of osteoblast lesion. Scale bar = 10 µm.

**Movie 5 (supplement to Fig. 4C):** Movie showing the recruitment of macrophages, labeled in magenta, into the area of osteoblast lesion. Scale bar = 10 µm.

**Movie 6 (supplement to Fig. 4E):** Movie showing the recruitment of macrophages from the surrounding tissue and not from blood vessels after osteoblast lesion. Scale bar = 20 µm.

**Movie 7 (supplement to Fig. 6):** Movie showing the impaired recruitment of macrophages, labeled in magenta, into the area of osteoblast lesion after pre-treatment with the antioxidant DPI. Scale bar = 10 µm.

**Movie 8 (supplement to Fig. 7B):** Movie showing recruitment of macrophages, labeled in magenta, into the area of osteoblast lesion after DMSO treatment. The response is comparable to the untreated response observed in laser-ablated, otherwise untreated zebrafish (see Movie 5). Scale bar = 10 µm.

**Movie 9 (supplement to Fig. 7B):** Movie showing reduced recruitment of macrophages, labeled in magenta, into the area of osteoblast lesion after prednisolone treatment. Please compare with the response in control treated (Movie 7) and untreated (see Movie 5) individuals. Scale bar = 10 µm.

**Movie 10 (supplement to Fig. 8B):** Movie showing recruitment of macrophages, labeled in magenta, into the area of a small osteoblast lesion after DMSO treatment. The response is weaker compared to the response observed in the larger laser-ablated osteoblast lesioned, otherwise untreated zebrafish (see Movie 5). Scale bar = 10 µm.

**Movie 11 (supplement to Fig. 8B):** Movie showing reduced recruitment of macrophages, labeled in magenta, into the area of a small osteoblast lesion after prednisolone treatment. Please compare with the response in control treated (Movie 9) and larger lesion prednisolone treated (see Movie 8) individuals. Scale bar = 10 µm.

